# Changing food conditions and size declines in a North Sea forage fish

**DOI:** 10.1101/2025.03.25.643521

**Authors:** Agnes B. Olin, Neil S. Banas, Michael R. Heath, Peter J. Wright, Alan MacDonald, Sarah Wanless, Francis Daunt, John R. Speakman, Ruedi G. Nager

## Abstract

Declining body sizes are prevalent in marine fish. While these declines have been suggested to be a response to increasing temperatures, the evidence is mixed and the underlying causes of observed declines often unknown. Here, we explore drivers of spatio-temporal patterns in size in lesser sandeel (*Ammodytes marinus*), an important prey for seabirds and marine mammals, focusing on ongoing size declines in the North Sea. We combine experimental and field data with ecological theory to develop a biologically realistic dynamic energy budget model that explicitly models feeding, metabolism and energy allocation to produce daily predictions of size during the juvenile growth season from 1979 to 2016. When forced with daily temperature estimates and zooplankton data from the Continuous Plankton Recorder, model predictions reproduce observed spatio-temporal patterns in size well. Our results suggest that the most plausible driver of observed size declines in the western North Sea is declining prey densities. In contrast, the direct effect of temperature on sandeel size is small, but interacts with local prey availability so that the effect varies in both size and direction over space. Our results thus suggest that to understand effects of climate change on fish size we need to account for both direct physiological effects and changes in resource availability. Finally, we use the model to show that early-life phenology and turbidity (via its impact on intake rates in the visually foraging sandeel) may also impact sandeel size, highlighting the importance of broadening our view of potential drivers of size declines.

## 1. INTRODUCTION

Declining body sizes have been proposed as a third “universal response” to climate change, in addition to poleward shifts in distribution and shifts in the timing of seasonal events (Daufresne et al. 2009, Gardner et al. 2011, Sheridan & Bickford 2011). However, the evidence to support this claim is mixed, where body sizes have been shown to both increase and decrease in response to warming (Teplitsky & Millien 2014, Audzijonyte et al. 2020) and trends towards increasing body sizes are equally common in most taxa (Martins et al. 2023). In contrast with other taxa, marine fish do, however, show consistent evidence of declining average body sizes (Martins et al. 2023). The drivers of these declines and how temperature increases may, or may not, affect fish body size is hotly debated. Several mechanisms have been invoked, such as faster development rates but smaller adult sizes (temperature-size rule), and increasing metabolic rates leaving less resources for growth at all ages (Gardner et al. 2011, Sheridan & Bickford 2011, Cheung et al. 2013, Ikpewe et al. 2021). Several other drivers, including declines in the abundance and quality of food (Korman et al. 2021, Menu et al. 2023, Queiros et al. 2024), size-selective predation and fishing (Swain et al. 2007, Ohlberger et al. 2019), and increased competition (Ohlberger et al. 2023) have also been proposed as key contributors to the size declines. In many cases, the drivers are not yet fully understood. However, teasing apart the underlying mechanisms is important, as size is strongly linked to survival (Levangie et al. 2022) and fecundity (Barneche et al. 2018) and thus affects both abundance and the quality of individual fish, with implications for both sustainable fisheries management (Audzijonyte et al. 2013, Persson et al. 2014) and the health of piscivorous predators (Österblom et al. 2001, Engelhard et al. 2014).

One species of forage fish that has exhibited pronounced declines in size is the lesser sandeel (*Ammodytes marinus*), a lipid-rich shoaling fish inhabiting sandy banks in the north-east Atlantic. It is an important trophic link between the zooplankton and several species of seabirds, marine mammals and piscivorous fish, as well as the target of a substantial fishery (Engelhard et al. 2014). The sandeel shows marked spatio-temporal variation in size-at-age in the North Sea region, with large and stable body sizes in the north-east, and smaller and declining sizes in the western North Sea (Bergstad et al. 2002, Harris & Wanless 2011, van Deurs et al. 2014, Rindorf et al. 2016, Wanless et al. 2018). The size declines in the western North Sea have been observed both in mature adults and in juveniles. Off the coast of southeast Scotland, declining juvenile body sizes from the mid-1970s to 2015 resulted in a 70% decline in energy content (Wanless et al. 2018), meaning that the sandeels’ predators had to catch over 3 times as many sandeels to fulfil their energetic needs.

The drivers behind the sandeel size declines are still unclear. Water temperatures, which are increasing rapidly in the north-east Atlantic (Kessler et al. 2022), have been linked to body size in both lesser sandeels and other *Ammodytes* species (Robards et al. 2002, Eliasen 2013, Rindorf et al. 2016). However, the direction of the relationship is inconsistent and modelling work suggests that temperature is not a strong driver of lesser sandeel growth (MacDonald et al. 2018). Variability in prey availability and composition has long been proposed as the main driver of spatial patterns in sandeel size (Macer 1966, Bergstad et al. 2002, Boulcott et al. 2007), with modelling work suggesting that food availability is a key driver of lesser sandeel growth (MacDonald et al. 2018). Prey availability has declined steeply in several of the locations where sandeel size has declined (Olin et al. 2022), which could have contributed to the observed temporal trends. A shift towards a later start to the growth season of juvenile sandeels has also been proposed as a potential driver of the size declines (Frederiksen et al. 2011), possibly driven by temperature-driven delays in spawning (Wright et al. 2017) and a mismatch between sandeel phenology and peak availability of larval food (Régnier et al. 2019, 2024). Finally, turbidity has increased within the sandeel’s range due to coastal erosion, intensified winds and waves resuspending more sediment, and bottom trawlers stirring up sediment and destroying beds of water-filtering bivalves (Capuzzo et al. 2015, Wilson & Heath 2019). As light conditions have been identified as a key driver of intake rates in the visually foraging sandeel (Winslade 1974b, van Deurs et al. 2015), this may therefore have contributed to the observed size declines. In contrast, neither competition (Rindorf et al. 2016, Henriksen et al. 2021) nor predation (Rindorf et al. 2016) or fishing (Bergstad et al. 2002, Wanless et al. 2004, Rindorf et al. 2016) appear strongly linked to lesser sandeel size.

This study aims to provide insight into causes of the marked declines in sandeel size, improving our understanding of drivers of size declines in fish in marine ecosystems under anthropogenic change. To do so, we use a dynamic energy budget model to explore drivers of growth in juvenile lesser sandeels in their first summer. Dynamic energy budget models track energy gains and losses as a function of environmental conditions (e.g. temperature, food) and then translate this into changes in body size and energy reserves (Kooijman 2000, Lika & Nisbet 2000). Such mechanistic models are helpful for teasing apart the roles played by different drivers, enabling us to gain better insight into the impact of ongoing environmental change on fish body sizes. The model builds on a dynamic energy budget model developed by MacDonald et al. (2018). However, the MacDonald model was parameterised specifically for the north-western North Sea and relies on several fitting parameters (e.g. prey type-specific attack rates, background food availability) that may vary over space (and time), requiring us to make adjustments in order to allow us to study the large-scale, long-term patterns we were interested in here. This involved breaking processes into tractable sub-processes that could be parameterised using data from experiments and measurements from the field, providing us with a biologically realistic model that can be more readily extended across space and time. We validate model predictions against field data and then use the model to explore to what degree observed spatio-temporal variation in juvenile sandeel size in the North Sea can be explained by the candidate drivers introduced above — (i) sea surface temperatures, (ii) food availability and composition, (iii) sandeel phenology and (iv) turbidity.

## 2. MATERIALS AND METHODS

### 2.1. Dynamic energy budget model

Here, we develop a dynamic energy budget model that covers the first growth season, from metamorphosis to winter dormancy, when the sandeels cease feeding and bury into the sand (MacDonald 2017, van Deurs et al. 2011b). As only a small proportion of sandeels spawn in their first year (<5 % in most areas; Boulcott et al. 2007), reproduction is not included in the model.

The model is constructed around two state variables: reserve energy *R* (kJ, remobilisable tissue, mostly fat) and structural energy *S* (kJ, non-remobilisable tissue, such as skeletal tissue). The basic structure involves the allocation of net energy gain (assimilated energy *A* [kJ day^-1^], minus metabolic costs *M* [kJ day^-1^]) to reserve energy and structural energy (see Figure 1). Assimilated energy is the energy from ingested food, after accounting for assimilation efficiency. The model assumes that metabolic costs are subtracted from assimilated energy and that if the assimilated energy is not enough to meet metabolic costs, the rest is subtracted from reserves. If the assimilated energy is larger than the metabolic cost, a certain proportion *f_S_* of this net energy gain is allocated to structural energy and the rest (1-*f_S_*) to reserve energy. Reserve energy *R* (kJ) thus changes as follows:

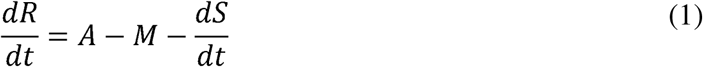

**Fig. 1.**
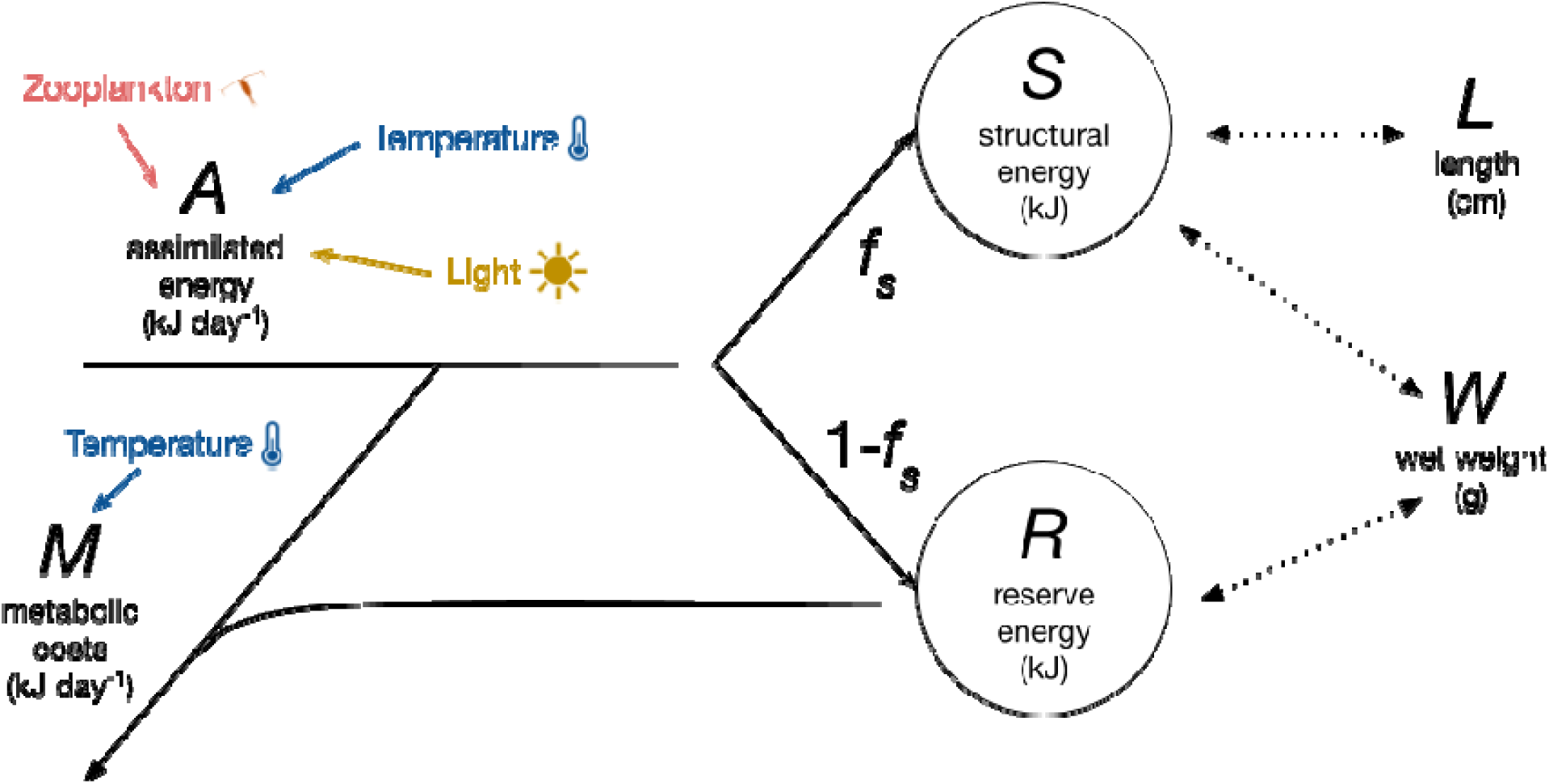
State variables and key processes in the dynamic energy budget model. Solid black arrows represent energy flows, coloured arrows environmental effects and dotted arrows the relationship between the state variables ( and ) and sandeel length and wet weight . is the proportion of net energy gain allocated to structural energy.

Structural energy *S* (kJ) then follows:

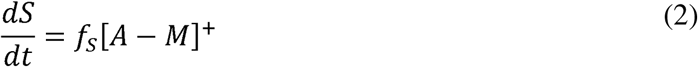

where [*A* – *M*]^+^ signifies that allocation to structural energy only occurs if net energy gain is positive. The equations are discretised assuming time steps of one day, thus providing daily estimates of reserve and structural energy. To be able to compare the model output to field observations, we translated reserve and structural energy into length and wet weight (see SI1).

The model is run from metamorphosis (generally mid-May, day 141, see “Initial conditions”) until early August (day 212), which is roughly when the growth season ends and the sandeels bury into the sand for winter (van Deurs et al. 2011b, MacDonald 2017). Each model component (ingestion, metabolism, energy allocation) is described briefly in the following subsections, with details provided in the SI (see also Olin 2020). Model parameters are presented in SI2 with descriptions of how the values were derived and an analysis of how sensitive model predictions are to choices of parameter values. The model is implemented in C, based on an adaptation of the growth component of the model presented in MacDonald et al. (2018). R 3.5.2 (R Core Team 2018) was used for data processing and visualisation.

#### 2.1.1. Assimilated energy

Prey availability and composition appear to be a major determinant of growth rates in lesser sandeels (van Deurs et al. 2014, 2015, MacDonald et al. 2018), and several studies indicate that sandeels feed selectively (Godiksen et al. 2006, Christensen 2010, Eliasen 2013). Therefore, particular attention was paid to modelling ingestion (see SI3 for details). This is where the main modifications to the MacDonald model were made, where sub-processes were isolated and modelled explicitly (see Olin 2020 for a detailed comparison of the models). Ingestion is modelled on an hourly basis, assuming a daily feeding window covering the hours of light (Freeman et al. 2004, Johnsen et al. 2017), minus one hour for school aggregation in the morning and one hour for school disintegration before the sandeels bury into the sediment for the night (see van Deurs et al. 2011a). Total assimilated energy per day *A* is then obtained by adding up the ingested energy for each hour of feeding, and multiplying it by the assimilation efficiency (proportion of energy remaining after faecal losses and nitrogenous excretion; Jobling 1993). Based on observations of other *Ammodytes* species, assimilation efficiency is assumed to increase linearly with temperature (Larimer 1992, Gilman 1994; SI3.1). Based on experimental observations, it is also assumed that the sandeels do not feed if there is not enough food to account for the metabolic costs of feeding (Winslade 1974a, van Deurs et al. 2011a). Total assimilated energy *A* (kJ day^-1^) is thus calculated as:

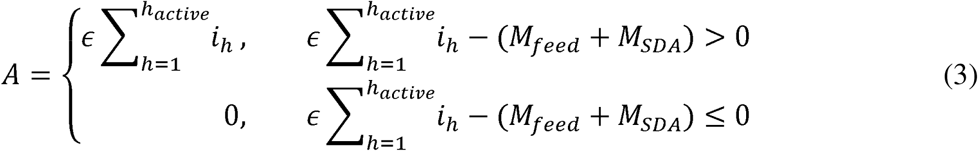

where *∈* is the assimilation efficiency, *h_active_* the number of hours feeding, *i_h_* the energy ingested during a given hour, and *M_feed_* and *M_SDA_* the cost of feeding and synthesising tissue, respectively (see below).

Hourly ingested energy *i_h_* is limited by the available prey as well as gut capacity. To incorporate this, we first modelled the maximum potential intake rate *i_max_* (kJ h^-1^; SI3.2) in response to the prey field, and then, if necessary, down-adjusted this according to remaining gut space (SI3.10). Gut content was therefore also modelled on an hourly basis, based on ingestion and digestion, the latter depending on both temperature and prey energy density (SI3.9). The response to the prey field was modelled as a Holling type II response (Holling 1959) into which we incorporated three forms of prey selectivity (Eggers 1977): (1) a prey size-, sandeel size- and light-dependent prey detection distance (SI3.4; this built on a sandeel foraging model by van Deurs et al. 2015), (2) a prey size-dependent capture probability (SI3.6) and (3) active switching (assuming switching behaviour is based on the profitability of each prey search class; see Visser & Fiksen 2013; SI3.2). All these forms of selectivity are supported by observations of sandeels (e.g. Godiksen et al. 2006, Christensen 2010, see Olin 2020 for details).

#### 2.1.2. Metabolism

The model includes three types of metabolic costs: (i) standard metabolic rate (SMR), which is the energy required to cover basic maintenance, (ii) costs associated with feeding behaviour and (iii) costs of synthesising tissue (specific dynamic action, SDA). Total metabolic costs *M* (kJ day^-1^) are calculated as:

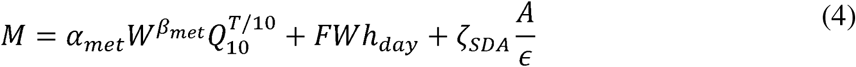

where *α_met_is* a constant, *w* is sandeel wet weight (g), *β_met_* is the weight-scaling exponent of SMR, *T* is temperature (°C), *Q*_10_ represents the relative change as temperature increases by 10°C, *F* is the foraging cost per hour per gram of sandeel, *h_day_* is the total number of hours spent out of the sand each day (thus assuming that the cost of school aggregation and disintegration is the same as the cost of foraging), *ζ_SDA_* is the SDA coefficient, *A* is the assimilated energy per day (kJ day^-1^) and *∈* the assimilation efficiency. It is thus assumed that SMR is a function of sandeel weight and temperature, the main predictors of SMR in fish (Clarke & Johnston 1999), that feeding costs are a function of activity and sandeel size, and that SDA is a function of the amount of ingested energy (see SI4 for details).

#### 2.1.3. Energy allocation

Each day, if the net assimilated energy (*A* – *M*) is positive, a proportion *f_S_* of this is allocated to structural energy (Eq. 2), and the rest to reserves (Eq. 1). Based on observations in *A. marinus* (Hislop et al. 1991) and other *Ammodytes* species (Sekiguchi et al. 1976, Robards et al. 1999, Danielsen et al. 2016), we assumed that allocation to structural energy decreases as the sandeel grows (see SI5). Further, as the lipid content of *A. marinus* increases rapidly after a winter of fasting (Hislop et al. 1991, Rindorf et al. 2016), we assumed that sandeels prioritise allocating energy to reserves when reserves are below a certain threshold (see SI5).

### 2.2. Locations

We ran the model in four locations (Figure 2): Dogger Bank (54.7°N 1.5°E), Firth of Forth (56.3°N 2°W), the East Central Grounds (hereafter: ECG; 57.6°N 4°E) and Shetland (59.8°N 1.3°W). The locations were chosen to represent a range of growth conditions, where the ECG is expected to show the fastest growth and Firth of Forth the slowest (Bergstad et al. 2002, Boulcott et al. 2007), and size declines have been reported in all locations apart from the ECG (Harris & Wanless 2011, van Deurs et al. 2014, Wanless et al. 2018). The locations represent distinct sub-populations and separate fisheries management areas, based on evidence from tagging, otolith microchemistry, larval drift modelling and genetic studies (ICES 2024).

**Fig. 2.**
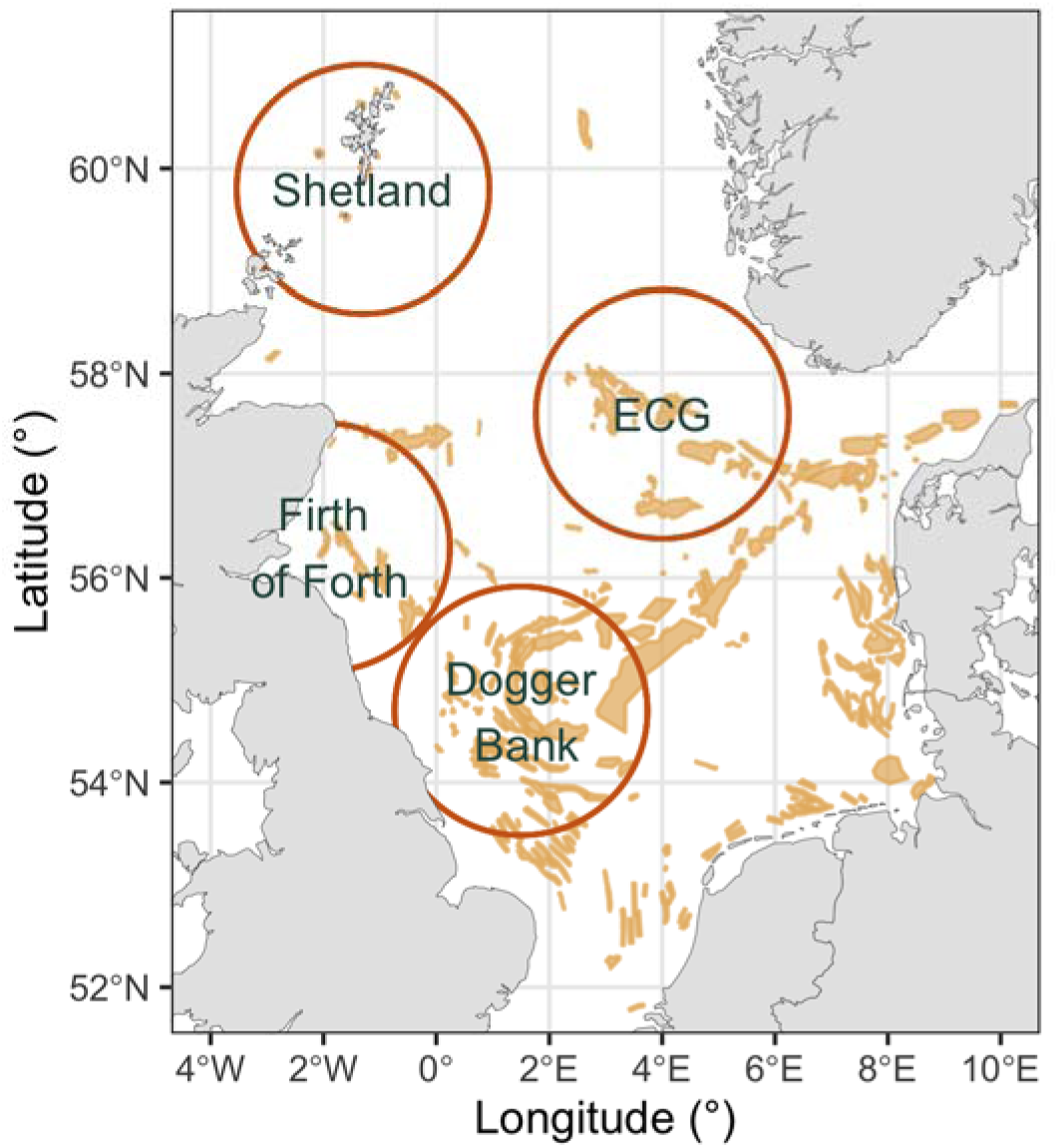
Study area, with each location marked with a circle indicating the area from which zooplankton data were sourced (see “Environmental data”). ECG = East Central Grounds. Shaded yellow areas indicate sandeel grounds (Jensen et al. 2011; data from Shetland from Marine Scotland Science).

### 2.3. Environmental drivers

The model requires the following environmental drivers: abundances, energy content, image area and length of each prey type, sea surface temperatures, day length, average surface solar irradiance and the diffuse attenuation coefficient *a_d_*, which depends on turbidity.

Daily prey abundances were based on data collected by the Continuous Plankton Recorder (see Olin et al. 2022 for methods). Based on prey found in sandeel stomachs, the prey taxa included copepods, Euphausiacea, Hyperiidea, Decapoda larvae, Appendicularia, fish eggs, fish larvae, *Evadne* spp. and *Podon* spp. A full list of prey taxa with energy content, prey image area, length and search class can be found in Table S2. The prey fields were based on data aggregated over a 135 km radius circle centred on each study location (see Olin et al. 2022; Figure 2). The chosen size of the area results from a trade-off between sample size and the homogeneity of the area it represents. The size of the area, and the between-sample variability in the alignment of zooplankton patches and the Continuous Plankton Recorder transects, means that the prey field input is not an exact representation of available prey in the study location for that year. Therefore, we would not necessarily expect the model to reproduce observed sandeel sizes in a given year, even if the model would correctly capture all relevant mechanisms. Instead, the model should be judged by its ability to capture long-term and large-scale spatio-temporal patterns.

We obtained temperature estimates from the ERA5 Climate Reanalysis, providing hourly sea surface temperature with a 31×31 km resolution (Copernicus Climate Change Service C3S 2017), averaged to daily values. As sandeels may forage throughout the water column and reside in hydrographically dynamic areas (Tien et al. 2017), it was assumed that surface temperatures were representative for the experienced temperatures at all depths. Hours of daylight were obtained using the function “daylength” in the R-package “geosphere” (Hijmans 2017). Average daily surface irradiance (SI3.5) was calculated using a Fortran subroutine (see Ljungström et al. 2020). The diffuse attenuation coefficient *a_d_* was obtained from observations in hydrodynamic regions corresponding to sandeel habitat (see supplementary materials in Capuzzo et al. 2018) and was assumed to be constant.

### 2.4. Initial conditions

The initial conditions of the model include size and day of year at metamorphosis. We used day 141 (21 May in a regular year) as the default starting date and 4 cm as the default starting length, chosen to be broadly representative for the study locations (Wright & Bailey 1996, Jensen 2000, Régnier et al. 2017).

### 2.5. Model validation

The model was run in all four locations for the years 1979–2016, excluding location-years in which insufficient zooplankton data (fewer than three samples per month) were available (Dogger Bank N = 33, Firth of Forth N = 23, ECG N = 23, Shetland N = 36). We then assessed whether the model could recreate observed large-scale and long-term spatio-temporal patterns in sandeel size, making use of all juvenile size observations we could locate from our study locations. This included (i) fisheries data from Shetland and the ECG collected in 1979 (Bergstad et al. 2002), (ii) dredge surveys in the Firth of Forth, Dogger Bank and a location slightly south of the ECG in 1999 (Boulcott et al. 2007), (iii) dredge surveys since 2006 in the ECG and since 2004 in Dogger Bank (ICES 2024), (iv) sandeels brought in by Atlantic puffins (*Fratercula arctica*) to the Isle of May in the Firth of Forth (Wanless et al. 2018) and (v) corresponding datasets of sandeels collected from puffins in the Shetland area, one from Fair Isle, south of Shetland, and one from Hermaness, in the north of Shetland (Harris & Wanless 2011). The first three datasets are representative of length at overwintering, while the latter two are standardised to the 1st of July. As the puffin dataset from the Firth of Forth was used to tune handling time to achieve the same mean size in the observations as in the predictions (see SI2), it is here only used to assess whether our predictions reproduce temporal trends. Temporal trends were assessed using linear regression.

### 2.6. Drivers of growth

The sensitivity of length predictions to our hypothesised drivers (temperature, food, phenology, light) was then investigated, quantified as the percentage difference in length at overwintering compared to a baseline scenario. To do this, the model was run for all location-years with data, varying one driver at a time while keeping the remaining input at their original values. This approach isolates the effect of individual drivers while also ensuring that the full range of environmental conditions are captured.

To examine the impact of temperature, a baseline annual cycle was established for each location by averaging the sea surface temperature for each day of the year across years. A range of temperature conditions were then examined by adjusting this baseline, from subtracting 3°C (corresponding to coldest year in dataset) to adding 4.5°C (similar to the temperature anomaly of the 2023 heatwave, Berthou et al. 2023). We also compared the average temperature over the growth season for a given year with (i) predicted lengths at overwintering, to assess the relative importance of temperature in driving model predictions, and (ii) actual observed lengths, to determine whether similar patterns are present in field data. As a humped relationship emerged when varying temperature across our defined range (see Results), this was done using both a simple linear regression and a second-order polynomial. To account for any temporal autocorrelation, the models were fitted with a first order auto-regressive error structure. The models were compared using ΔAIC_C_.

To investigate the role of food, we focused on three aspects: the total amount of available energy, the density of *Calanus finmarchicus* (often identified as a key driver of bottom-up dynamics in this region; Frederiksen et al. 2013, van Deurs et al. 2014) and the prey size, where the availability of large prey is thought to boost sandeel ingestion and growth rates (van Deurs et al. 2015, MacDonald et al. 2018). First, for each location, we varied the total amount of energy available throughout the whole season from the lowest to the highest observed value in the time series by applying a constant scalar to daily zooplankton densities, thus maintaining seasonal patterns and keeping the relative density of each taxa constant. Then, we repeated this approach but instead varied only the density of *C. finmarchicus*, from the lowest to the highest mean density observed in each location, keeping all other prey types at their original densities. Again, the seasonal pattern was preserved. For the prey size we took a different approach, exploring the effect of keeping the total available energy for a given day unchanged, but having all energy in just one prey type. The prey types we explored included *Oithona* spp. (0.68 mm), *Acartia* spp. (1.15 mm) and *C. finmarchicus* (2.7 mm), considered representative of small, medium and large prey, respectively. As for temperature, we compared the daily energy availability, average daily *C. finmarchicus* densities and average prey size over the growth season for a given year with (i) predicted lengths at overwintering and (ii) actual observed lengths. This was done using both a simple linear regression and a log_10_-transformation, as the positive effects were expected to level out. Again, the models were fitted with a first order auto-regressive error structure and were compared using ΔAIC_C_. For (ii), we note again that the representativeness of prey field data may vary between years, so results should be interpreted with caution.

To assess the impact of phenology and larval growth processes on predicted size, the impact of date of metamorphosis and size at metamorphosis was examined. The model start date (metamorphosis date) was varied from 121 to 181, and the initial size (metamorphosis length) was varied from 3.5 to 5.5 cm based on observed ranges (Wright & Bailey 1996, Jensen 2000, Régnier et al. 2017, 2024). As the prey field input was kept constant, varying the model start date is equivalent to examining the role of variation in sandeel phenology relative to prey phenology.

Finally, to examine the impact of light conditions, the diffuse attenuation coefficient *a_d_* was varied over the range 0 (completely clear waters) to 0.3, based on a range of values commonly observed in the type of hydrodynamic region corresponding to sandeel habitat (see supplementary materials in Capuzzo et al. 2018).

## 3. RESULTS

### 3.1. Model validation

While tuned only to length data from the Firth of Forth, the model also produced realistic predictions for the other locations and reproduced spatial differences in size (Figure 3). Both observations and predictions suggest that (i) in the late 1970s, growth conditions in the ECG were better than in Shetland, (ii) in the late 1990s, growth conditions were better in the ECG than in Dogger Bank, which in turn were better than in the Firth of Forth, and (iii) the better growth conditions in the ECG compared to the Dogger Bank were maintained in the 2000s and 2010s (Figure 3a–d).

**Fig. 3.**
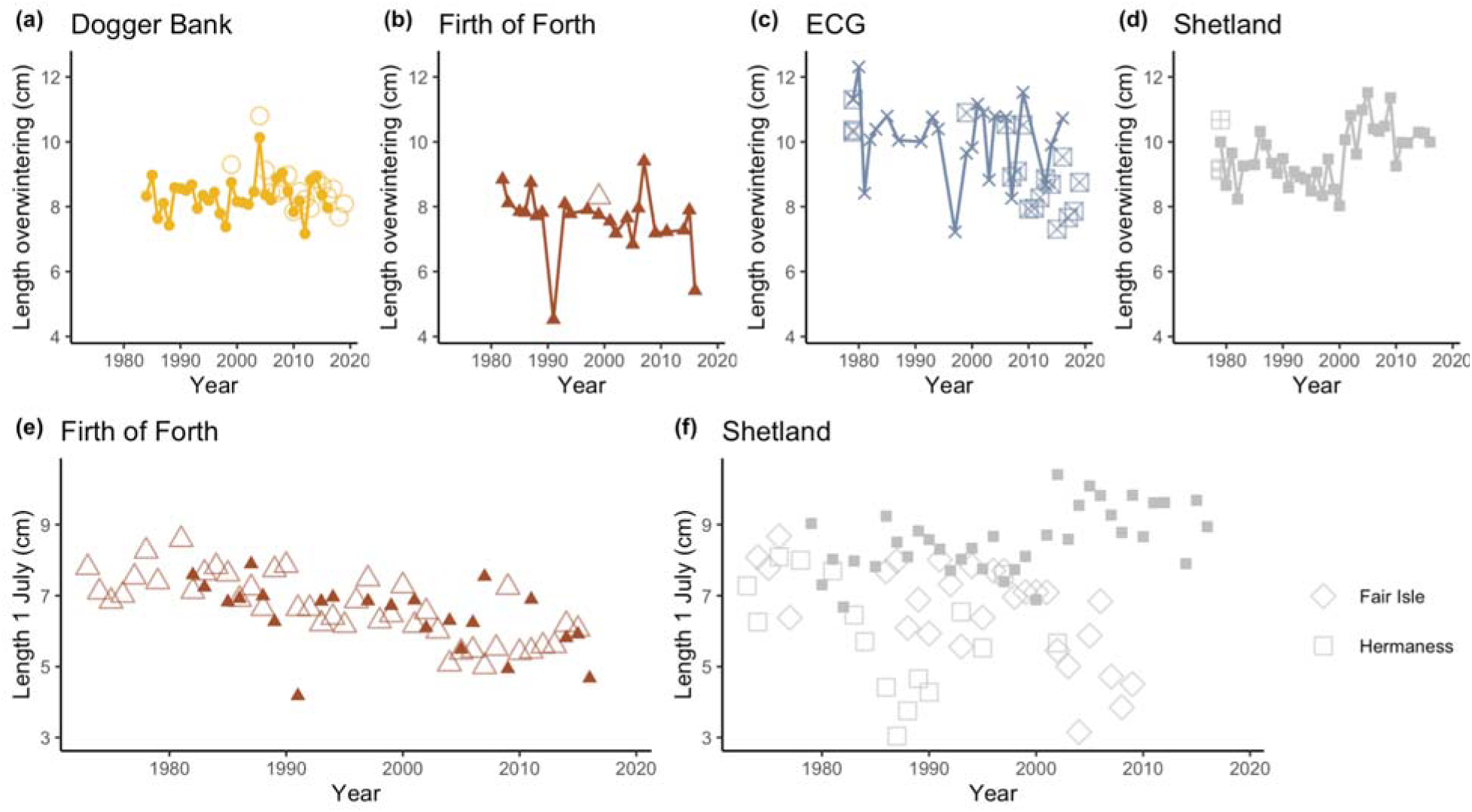
(a–d) predicted lengths at overwintering for (a) Dogger Bank, (b) Firth of Forth, (c) ECG (East Central Grounds) and (d) Shetland, with corresponding observational data as described in the Methods (large open symbols). (e–f) predicted lengths on the 1st of July for (e) Firth of Forth and (f) Shetland, with corresponding observational data as described in the Methods (large open symbols).

The model predictions also did well in reproducing the temporal trend in the Firth of Forth (Figure 3e). Observations showed a decline in sandeel size between 1982 and 2015 of -0.06 [95 % CI: -0.08; -0.04] cm per year. Predictions over the same time period also showed some evidence of a decline, and although a weaker decline of -0.03 [95 % CI: -0.07; 0] cm per year, the 95 % confidence intervals of the two slopes overlapped. In Shetland, predictions pointed to an increase in size by 0.05 [95 % CI: 0.02; 0.09] cm per year (Figure 3f) over the time period 1979–2009 (the years for which we had both predictions and observations). This does not align with the observations from Fair Isle, which instead showed a decline of -0.12 [95 % CI: -0.19; -0.06] cm per year, or from Hermaness, where no trend was observed [estimate: 0.02; 95 % CI: -0.15; 0.19].

In Dogger Bank for the period 2004–2016 when both predictions and observations are available, neither observations [estimate: 0; 95 % CI: -0.08; 0.07] nor predictions [estimate: -0.03; 95 % CI: -0.16; 0.09] showed any trend. There was also no trend in predicted size over the time period 1988–2011 during which size declines have been reported in older age groups (van Deurs et al. 2014) [estimate: 0.02; 95 % CI: -0.02; 0.05]. In the ECG for the period 2006–2016 when predictions and observations are both available, neither observations [estimate: -0.14; 95 % CI: -0.35; 0.07] nor predictions [estimate: 0.01; 95 % CI: -0.43; 0.45] showed any trend.

### 3.2. Drivers of growth

#### 3.2.1. Temperature

Varying the temperature had a minor impact on predicted sandeel length (<1 % compared to baseline, Figure 4a). The effect was nonlinear, with increased temperatures resulting in increased growth up to an optimum after which growth instead decreased. The location of the optima in relation to the baseline varied between locations, so that a temperature increase would likely result in a small decline in body size in the Firth of Forth and Dogger Bank (optima just below average temperatures over the study period), whereas boosted growth was predicted for the ECG and Shetland (optima close to maximal warming). There were no relationships between observed growth season temperatures and predicted length at overwintering (Table S3; Figure 4b). However, in Fair Isle, we saw a negative linear relationship between growth season temperatures and actual observed lengths, where length decreased by 1.6 [95 % CI: 0.92; 2.2] cm per 1°C increase (Table S3; Figure 4c).

**Fig. 4.**
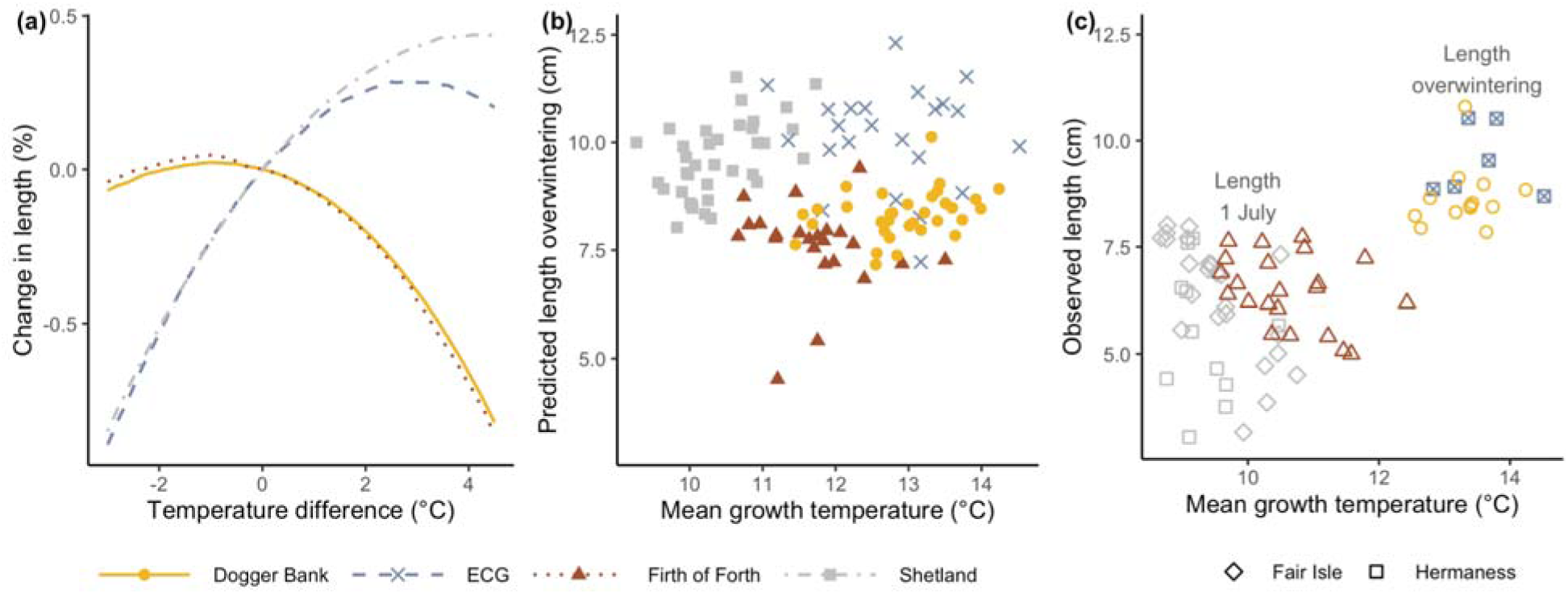
Effect of temperature on sandeel length. (a) effect of temperature on predicted lengths at overwintering, in relation to predictions at average temperatures (temperature difference = 0°C). (b) average temperature across the growth season compared to predicted length at overwintering. (c) average temperature from metamorphosis until date of length observations against observed length from field data. Note that for (c), the date of observation varies between locations so that values cannot be compared across locations. Closed symbols are predicted length (b), open symbols observed length (c). ECG = East Central Grounds.

#### 3.2.2. Food

Predicted growth was sensitive to average daily energy availability, where a shift from mean to maximum values resulted in a predicted increase in length of up to 14 % and a shift from mean to minimum values resulted in a predicted decrease of up to 38 % (Figure 5a). There were positive, log-shaped relationships between observed average daily energy availability in a given year and predicted length in the same year in the ECG and in Shetland (Table S4; Figure 5b). There was a positive, log-shaped relationship between observed average daily energy availability in a given year and observed length in the same year in Dogger Bank (Table S4; Figure 5c).

**Fig. 5.**
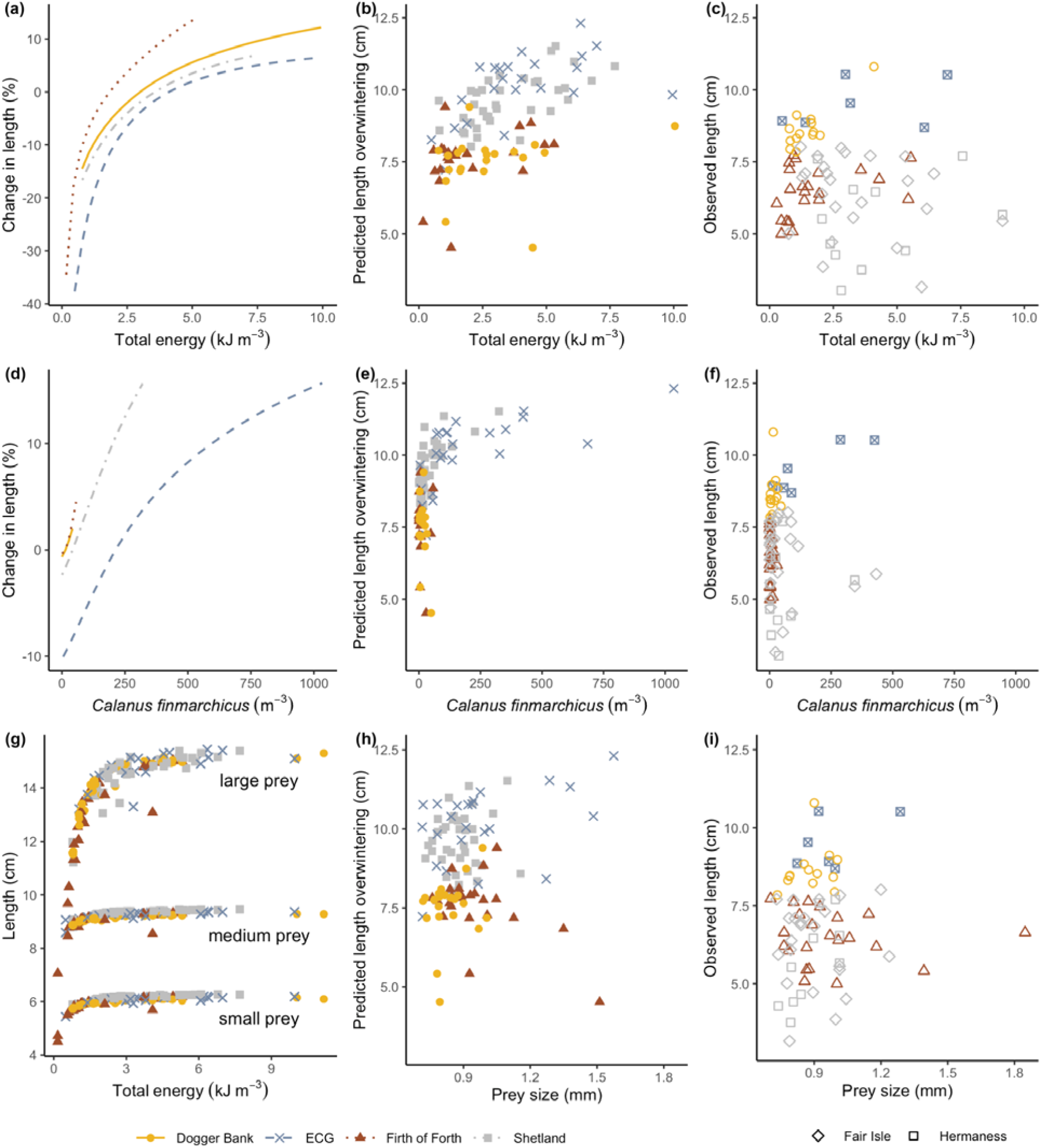
Effect of food conditions on sandeel growth. (a;d;g) effect of (a) average daily energy availability, (d) density of *Calanus finmarchicus* and (g) prey type on predicted lengths at overwintering. For (a) and (d) length predictions were averaged across years, and predictions ar presented in relation to average values. For (g) total available energy for a given day was kept unchanged, but all energy was provided in the form of large, medium, or small prey. (b;e;h) predicted length at overwintering compared to (b) average daily energy availability, (e) density of *Calanus finmarchicus* and (h) average prey size across the growth season. (c;f;i) actual length observations from field data compared with (c) average daily energy availability, (f) density of *Calanus finmarchicus* and (i) average prey size across the growth season from metamorphosis until date of length observations. Note that for (c;f;i), the date of observation varies between locations so that values cannot be compared across locations. Closed symbols are predicted length (b;e;h) and open symbols observed length (c;f;i).

For *Calanus finmarchicus*, there were clear differences between locations in the role it played. In the Firth of Forth and Dogger Bank, shifting densities over the range observed only resulted in a change in predicted length of ca. 1–5 %, whereas in the ECG, a shift from mean to maximum values resulted in a predicted increase of 16 % and a shift from mean to minimum values resulted in a predicted decrease of 10 %, and the corresponding values for Shetland were 16 % and 2 %, respectively (Figure 5d). There were positive, log-shaped relationships between observed *C. finmarchicus* densities and predicted length in the ECG and in Shetland (Table S4; Figure 5e). We saw no relationships between observed *C. finmarchicus* densities and observed length (Table S4; Figure 5f).

Prey type had a large effect on predicted lengths. For all three prey size classes examined, the predicted lengths increased with total available energy, but at peak energy availability, the predicted length for sandeels was ca. 15 cm when prey was supplied as large *C. finmarchicus*, whereas it was only ca. 6 cm when prey was supplied as small *Oithona* spp. (Figure 5g). There was a positive, linear relationship between observed average prey size during the growth season and predicted length in the ECG, and a negative, linear relationship in the Firth of Forth (Table S4; Figure 5h). In Hermaness, there was a positive, linear relationship between average prey size and observed length (Table S4; Figure 5i).

#### 3.2.3. Timing and size at metamorphosis

The effect of timing of metamorphosis was larger than the effect of size at metamorphosis (Figure 6). For the nominal value of size at metamorphosis (4 cm), a shift to the earliest date examined (day 121) resulted in a predicted increase in length at overwintering of 4–7 %, whereas a shift to the latest date examined (day 181) resulted in a decrease of 10–23 %. For the nominal value of timing of metamorphosis (day 141), a shift to the smallest size examined (3.5 cm) resulted in a predicted decrease in length at overwintering of 1–2 %, whereas a shift to the largest examined (5.5 cm) resulted in an increase of 3–7 %.

**Fig. 6.**
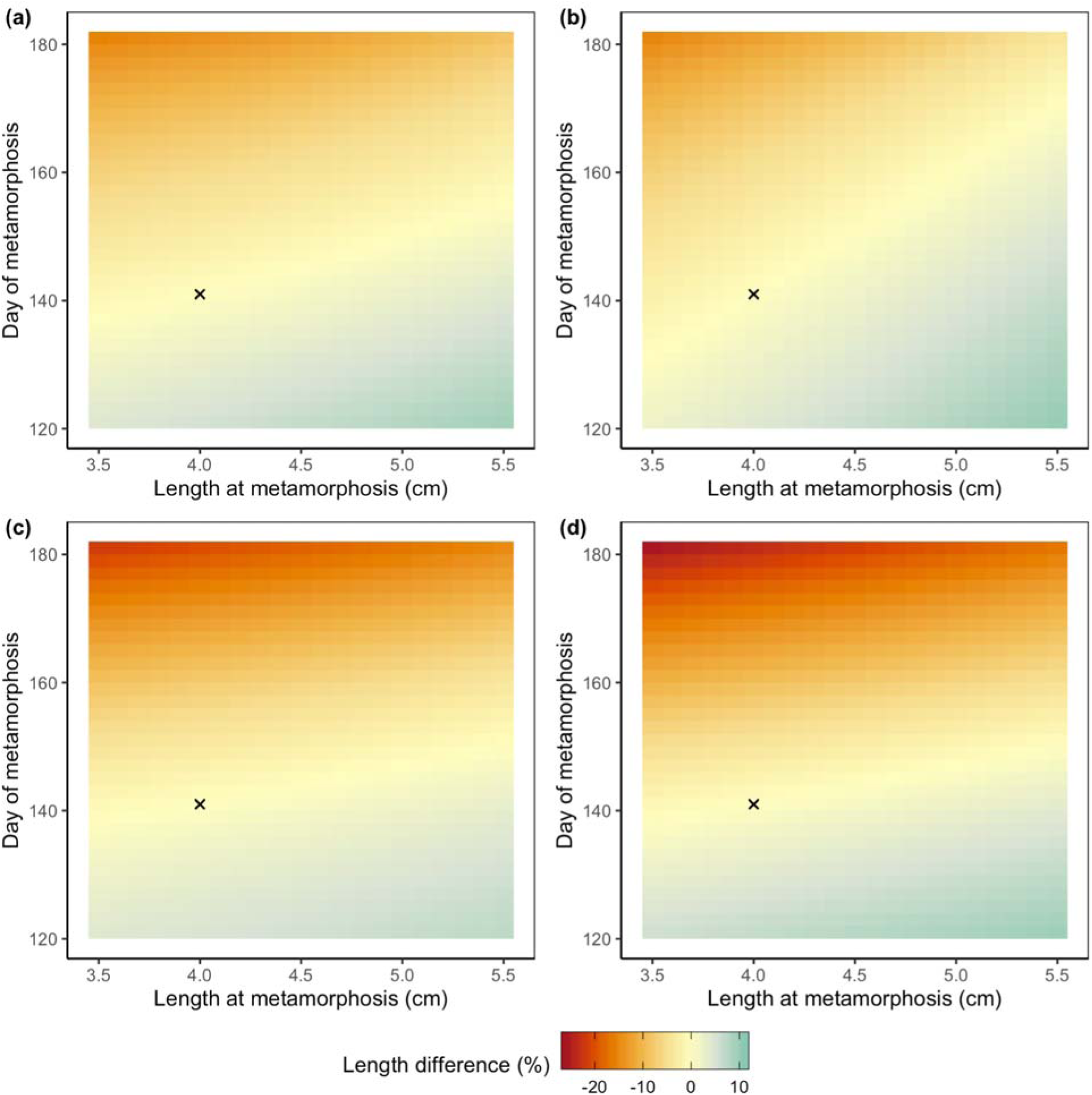
Effect of size at and timing of metamorphosis on predicted lengths at overwintering, in relation to predictions for nominal values (marked with x) for (a) Dogger Bank, (b) Firth of Forth, (c) ECG (East Central Grounds) and (d) Shetland.

#### 3.2.4. Light conditions

A shift towards increased turbidity (higher values for the diffuse attenuation coefficient *a_d_*) resulted in a decline in predicted sandeel size of up to ca. 50–60 % (Figure 7). A shift to completely clear waters only increased predicted length with ca. 3 %.

**Fig. 7.**
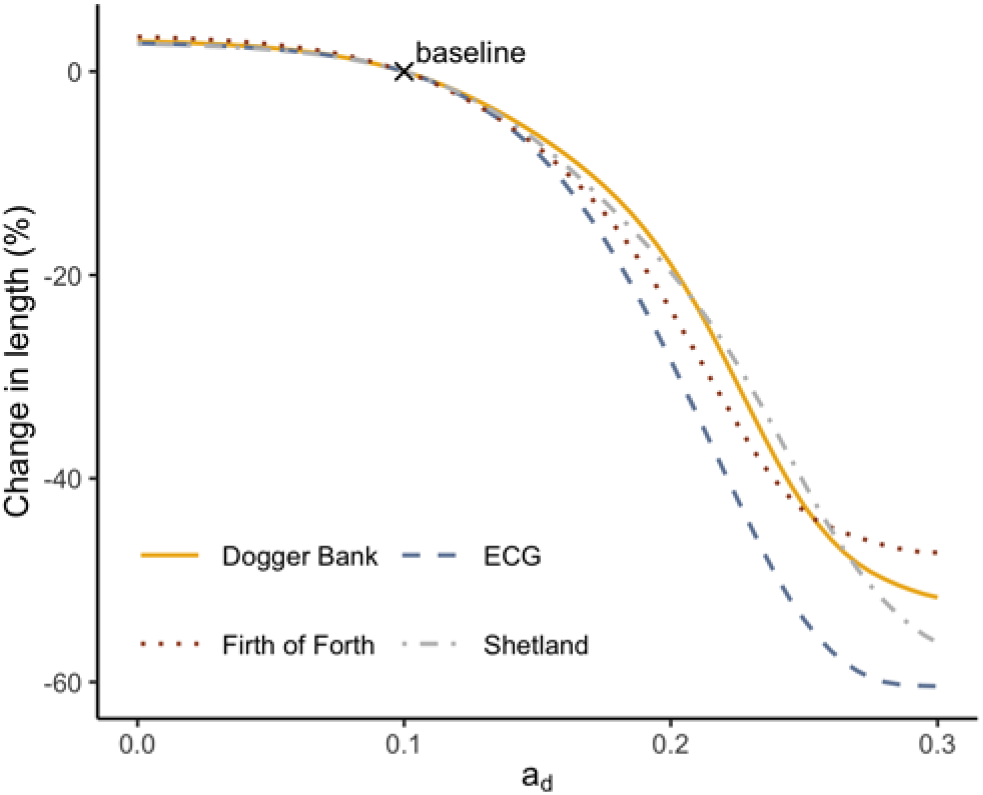
Effect of turbidity on predicted lengths at overwintering, in relation to predictions for the nominal value ( = 0.1). ECG = East Central Grounds.

## 4. DISCUSSION

This study used a dynamic energy budget model to explore plausible drivers of spatio-temporal variation in the growth of lesser sandeels in the North Sea region, with a particular focus on observed size declines. Model predictions matched observed spatio-temporal patterns well. Our results suggest that the effect of temperature on sandeel growth was minor, but varies in direction over space, and that it is unlikely that direct effects of increasing temperatures explain the size declines. In contrast, our results indicate that composition and density of prey are important drivers of sandeel growth rates. Variation in the timing of metamorphosis, and thus the start of the growth season, may also play a role in driving variation in size. Finally, turbidity could potentially have a large impact on sandeel growth via its effect on prey detectability.

As the direct effect of temperature was small, and light conditions as well as size at and timing of metamorphosis were kept constant, the model’s ability to reproduce the decline in size observed in the Firth of Forth suggests that trends in the composition and abundance of prey were sufficient to explain the observed size decline. This supports the hypothesis that a change in food conditions may be one of the key mechanisms behind the widespread declines in size observed in many organisms (Gardner et al. 2011), including fish (Korman et al. 2021, Menu et al. 2023). The decline in available food for the sandeels is primarily driven by declining abundances of small copepods (Olin et al. 2022). This explains the negative correlation between prey size and predicted sandeel length in the Firth of Forth (Figure 5h), as the larger average prey sizes result from low densities of small copepods (Olin et al. 2022). Tyldesley et al. (2024) showed that these declines in small copepod densities, and in total energy available to planktivorous fish, are widespread across the northwest European shelf and beyond, extending as far as Iceland and the southern Bay of Biscay. The ultimate driver is not known, but it could be linked to a decline in local primary productivity associated with increasing temperatures and decreased nutrient input (Capuzzo et al. 2018), possibly together with reduced quality and shifting phenology of the phytoplankton (Schmidt et al. 2020). The decline in sandeel size could thus still be related to climate change, via a change in prey availability and composition.

Food conditions are also a plausible driver of the spatial patterns in sandeel size, as long hypothesised (Macer 1966, Bergstad et al. 2002; Boulcott et al. 2007). Densities of *Calanus* spp. are higher in the north (Olin et al. 2022), and correlate with predicted growth in the ECG and Shetland (Figure 5e). Our results further suggest that a prey field composed of large, *Calanus*-like prey provides better growth conditions than smaller prey types, even when the total amount of energy is the same (Figure 5g), corroborating previous work showing the importance of prey size, in both lesser sandeels and other species (van Deurs et al. 2015, MacDonald et al. 2018, Ljungström et al. 2020). It is thus likely that the high densities of *Calanus* spp. explain why the sandeels grow so fast in the ECG. Previously, *C. finmarchicus* dominated the study area, but since the early 2000s, they are increasingly being replaced by *C. helgolandicus* (Olin et al. 2022, Tyldesley et al. 2024). This is likely the result of an ongoing temperature-driven northward distribution shift of both species (Edwards et al. 2020). In the northern North Sea, this shift may have a negative effect on sandeel growth in the long term, as the phenology of *C. helgolandicus* is less well matched with the sandeel growth season, and as peak densities of *C. helgolandicus* are unlikely to match those of *C. finmarchicus* (see Edwards et al. 2020, Olin et al. 2022). In comparison, densities of *C. finmarchicus* are, and have been, lower in the western North Sea, and are not positively correlated with predicted or observed sandeel growth in this area (Figure 5e–f). This is in line with recent evidence from Dogger Bank (Henriksen et al. 2018) and the Firth of Forth (Régnier et al. 2017, MacDonald et al. 2018) suggesting that the role of *C. finmarchicus* in driving sandeel dynamics may have been exaggerated in these areas (see e.g. van Deurs et al. 2009, 2014, Frederiksen et al. 2013).

The direct effect of temperature on sandeel growth was minor, resulting in a 1% difference in predicted size at most, even at a 4.5°C increase (Figure 4a). While our approach did rely on the assumption that sea surface temperatures are indicative of temperatures throughout the water column, this would, if anything, have led to an overestimated effect as variability is likely smaller deeper down. The small direct effect of temperature is in line with results presented by MacDonald et al. (2018), which were also based on a dynamic energy budget model of lesser sandeel, as well as work on other fish species (e.g. Menu et al. 2023). This suggests that temperature-driven increases in metabolic costs, which have been proposed as one of the mechanisms behind climate change-associated body size declines (Sheridan & Bickford 2011), are not the main cause of sandeel size declines.

In our model, warmer temperatures lead to greater assimilation efficiency and faster digestion rates, which allows for higher intake and growth rates. However, warmer temperatures also lead to increased metabolic costs, which result in a negative net energy gain if the increased costs are not outweighed by increased energy assimilation, which may be the case if food conditions are poor. This is why our study locations responded differently to temperature, where warmer temperatures led to higher growth rates in locations where food conditions are good (ECG, Shetland, high densities of *Calanus* spp.), but reduced growth rates where food conditions are poorer (Firth of Forth, Dogger Bank). This means that if a changing climate results in poorer food conditions, declines in growth rates may be mildly exacerbated by the increased metabolic costs of higher temperatures. Régnier et al. (2024) identified a similar pattern in sandeel larvae, where temperature had a positive effect on growth when the match between sandeel hatching and the peak availability of larval food was good, while the effect was instead negative when the match was poor. This type of interaction has also been noted in other fish species (Brett et al. 1969, Allen & Wootton 1982, Ohlberger 2013). Our study supports the claim that to understand effects of climate change on fish, we need to account for both direct physiological effects and changes in resource availability (Huey & Kingsolver 2019, Lindmark et al. 2022).

Our model only covered the first growth season, and therefore we could not fully evaluate the support for the temperature-size rule (i.e. fast development and smaller size-at-maturation, e.g. Gardner et al. 2011, Ikpewe et al. 2021). However, our model did suggest that if food is sufficient, temperature increases do result in boosted juvenile growth, in line with the temperature-size rule. As for size-at-maturation, a study from 1999 showed that Firth of Forth sandeels matured at a smaller size than Dogger Bank sandeels, which in turn matured at a smaller size than ECG sandeels (Boulcott et al. 2007). As the average annual temperature in the Firth of Forth was lower than in the other two locations, this does not fit with a smaller size-at-maturation in warmer temperatures. However, a more recent study found no significant differences in the relationship between size and maturation rates in these locations (Wright et al. 2019), and the difference in average annual temperatures between the locations is small (usually <1°C), so a broader geographical area would likely be needed to evaluate the support for the temperature-size rule in lesser sandeels.

While the potential effect of timing of metamorphosis on sandeel length-at-age was considerable, it cannot explain the size declines in the Firth of Forth on its own: the model predicts that a shift from the earliest to the latest observed date of metamorphosis in the Firth of Forth (Régnier et al. 2017) only results in a length difference of around 12% and thus cannot alone explain the decline in length of 28% over the study period. Further, there is no marked temporal trend (or spatial pattern) in larval or settlement phenology within the study area (Lynam et al. 2013, Régnier et al. 2019, 2024), and estimates of date at settlement for the Firth of Forth from recent years actually suggest that they are rather in the earlier part of the range we examined (Régnier et al. 2024). This provides further support for deteriorating food conditions as the most plausible driver of observed size declines in the Firth of Forth. Still, considering that phenology shifts are a common response to climate change in marine ecosystems (Poloczanska et al. 2013), temporal mismatch with prey may be a useful driver to consider in other cases of marine fish size declines.

As for the effect of turbidity, the potential role played was large. The findings here echo those based on studies of visually foraging fish in general (Aksnes 2007, Ljungström et al. 2020, Korman et al. 2021) and of *A. marinus* in particular (van Deurs et al. 2015). Turbidity in the North Sea varies seasonally and over space (e.g. Capuzzo et al. 2013) and has increased over time (Capuzzo et al. 2015, Wilson & Heath 2019). While satellite- and model-based estimates of turbidity are available for the North Sea, they do not extend far enough back in time to use them as input for the model. An interesting avenue for future research would be to explore spatio-temporal patterns in turbidity in sandeel grounds using these datasets. Increasing turbidity may also impact the foraging success of visually foraging sandeel predators (Finney et al. 1999, Lewis et al. 2015, Darby et al. 2022), suggesting that impacts could amplify up the trophic chain.

While the model predictions generally agreed with observations, this was not the case in Shetland. However, while predicted lengths were greater than those of the sandeels collected by puffins, they do match observations from trawl surveys from 1990–1992 (Wright & Bailey 1996; see Figure 4.2 in Olin, 2020) and from 2002–2007 (Marine Scotland Science, unpubl. data), the latter estimating mean juvenile lengths in August to 8–10.5 cm, overlapping with predictions at overwintering for the same time period (9.6–11.5 cm). As the time series from Fair Isle and Hermaness show different trends, possibly since the Fair Isle puffins go south to Orkney to forage, it is also difficult to know whether the size of Shetland sandeels have changed over time. Still, there is no empirical data that support the predicted increase, which is driven by an increased availability of food (Olin et al. 2022). Possibly, the benefits of increasing food availability during the juvenile feeding season have been cancelled out by a decline in food availability for fish larvae in the early 2000s (see Alvarez-Fernandez et al. 2012). Interestingly, poor breeding success and delayed breeding of sandeel-eating seabirds was also observed in the early 2000s in this region (JNCC 2016, Maniszewska 2019, Olin et al. 2020) suggesting that this time period warrants further study. The negative relationship between observed length and growth season temperatures in Fair Isle may also be worth exploring further.

In Dogger Bank, the predictions did not show evidence of a decline over the time period during which size declines have been reported in age 1 and age 2 sandeels (1988–2011; van Deurs et al. 2014). However, a closer examination of the published time series suggests that a significant decline only occurred in age 2 sandeel (see Figure S3 in SI6). No data exist on juvenile sandeel from the same time period so it is unclear whether a size decline in juvenile sandeels has actually occurred in Dogger Bank. Over the time period for which we have both observations and predictions, no decline was evident (Figure 3a). A difference in trends between age groups may imply that additional mechanisms are at play in older, mature sandeels, for example increased investment into reproduction as temperatures increase (see Wootton et al. 2022).

In summary, our results suggest that if we continue on the current trajectory of increasing temperatures (Kessler et al. 2022) prompting delays in phenology (see Wright et al. 2017, Régnier et al. 2019), increasing turbidity (Capuzzo et al. 2015, Wilson & Heath 2019), as well as shifts from *C. finmarchicus* to *C. helgolandicus* in the northernmost areas and declining densities of small copepods in the southernmost areas (Edwards et al. 2020), sandeel sizes may decline further. Since the possible drivers identified in this study are difficult to manage at a local or regional scale, another option to safeguard prey resources for marine top predators in the future may be to reduce fishing pressure on sandeels. Our results suggest that sandeel growth conditions have deteriorated in the western North Sea, which have long been important sandeel fishing grounds, and as smaller sandeels have higher mortality rates, lower maturation rates and lower fecundity (Boulcott et al. 2007, Boulcott & Wright 2011, MacDonald et al. 2018) this may make the sandeel stock vulnerable to additional mortality from fishing. A complete closure of sandeel fisheries in Scottish waters and in English waters within the North Sea (including Dogger Bank) were implemented in 2024, although it is currently unclear whether it is compatible with agreements with the EU (EU 2024).

The study provides some lessons of general interest. First, it lends support to the idea of temperature not as a driver that directly and uniformly pushes fish towards smaller body sizes, but rather a driver with complex direct and indirect effects, which may ultimately also result in increases in size in some contexts (see also Audzijonyte et al. 2020). Second, it highlights the importance of considering the prey field from the point of view of the predator, with a local perspective. It is tempting to identify key metrics such as, for example, the abundance of key prey taxa or average size of the prey to try to explain variation in growth. However, due to complex interactions depending on both prey and predator size and acting via, for example, capture success and switching mechanisms, the relationship between these metrics and predator growth may not always play out in a linear fashion, and may break down when extrapolating across space. For example, variation in *C. finmarchicus* densities is a good predictor of growth in the northern North Sea, but not in the western North Sea. Resolving these foraging dynamics may thus improve our understanding of how oceanographic change travels up the food chain all the way to top predators. Finally, the results also highlight the importance of broadening our view when it comes to identifying drivers of size declines. Our oceans are not only becoming warmer, but there are also trends, in various directions, in top predator densities, nutrient levels and fishing pressure, just to mention a few examples. A broader view of potential drivers helps to better partition variation between different mechanisms, ultimately improving our understanding of how marine ecosystems are responding to an increasingly changing environment.

## Acknowledgements

We want to thank Anna Rindorf, Dougie Speirs, Mikael van Deurs, Quentin Quieros and Max Lindmark for valuable input. We are also grateful to Espen Johnsen and Mikael van Deurs for providing sandeel data. A.B.O. was funded by a doctoral fellowship from the Marine Alliance for Science and Technology Scotland (MASTS), in partnership with the University of Strathclyde and the University of Glasgow. The present work was part of A.B.O.’s PhD thesis. N.S.B. was supported by the UK Missing Salmon Alliance under the Likely Suspects Framework project and the Natural Environment Research Council (Grant Number NE/X008983/1). R.G.N. was supported by the Natural Environment Research Council and the Department for Environment, Food, and Rural Affairs (Grant Number NE/L003090/1). We thank Mike Harris, Mark Newell and all members of the Isle of May field teams for the collection of puffin prey data. The Isle of May study was funded by NERC Award number NE/R016429/1 as part of the UK-SCaPE programme delivering National Capability.

