## Supplementary Information for "Changing food conditions and size declines in a North Sea forage fish"

This supplementary material contains further details and additional model equations for the lesser sandeel dynamic energy budget model. SI1 provides further information on the process of translating between length/weight and structural energy/reserve energy. SI2 includes a table with all parameter values as well as the result from a sensitivity analysis. SI3, SI4 and SI5 details the equations regarding assimilated energy, metabolic rate and energy allocation, respectively. See Olin (2020) for an extended description of the model. SI6 contains additional results.

### SI1: Translation

Structural energy $S$ (kJ) is directly related to length $L$ by a power function (see Broekhuizen et al., 1994) as follows:

$$S =\alpha_{dry}L^{\beta_{dry}}\delta_{S}$$

(S1)

where $\alpha_{dry}$and $\beta_{dry}$ are constants governing the translation of length into structural dry weight (g), $L$ is sandeel length (cm) and $\delta_{S}$ is structural energy density (kJ g dry weight^-1^).

Reserve energy $R$ (kJ) is instead a function of both length and wet weight:

$$R =\delta_{R}\frac{W-{\omega_{SDW}\alpha}_{dry}L^{\beta_{dry}}}{\omega_{RDW}}$$

(S2)

where $L$ (cm) is sandeel length, $W$ (g) is sandeel wet weight, $\omega_{SDW}$ and $\omega_{RDW}$ are constants for translating dry weights into wet weights for $S$ and $R$, respectively, $\alpha_{dry}$and $\beta_{dry}$ are constants governing the translation of length into structural dry weight (g) and $\delta_{R}$ is reserve energy density (kJ g dry weight^-1^).

See Table S1 for the estimation of parameter values. The defined relationships and estimated parameter values were tested against independent validation data, achieving good agreement (see Olin, 2020). Our approach to modelling energy allocation was similar to that of MacDonald et al. (2018) and parameter values are also similar. However, some of the parameter values were updated based on a detailed dataset on *A. marinus* body composition so that all values were derived from *A. marinus* data, which was not the case in MacDonald’s model.

### SI2: Parameter values

All variables and parameters are presented in Table S1, together with information on how the parameter values were derived. For three parameters (effective handling time $h_{e}$, and the two parameters governing capture probability as a function of prey size, $b$ and $m$), there were no previous studies or data that could be used to inform these parameters and these parameters were thus manually tuned (see Olin, 2020 for details). $b$ and $m$ were tuned first, using values from Pepin et al. (1987) as a starting point, and tuning the value to align with observed ratios between size of ingested prey and size of available prey, based on data from Godiksen et al. (2006). Effective handling time $h_{e}$ was tuned against a time series of sandeel lengths from the Firth of Forth, so that mean predicted and observed lengths across years were equal. This time series is based on sandeels brought in by Atlantic puffins (*Fratercula arctica*) to the Isle of May (56.2°N 2.6°W) in the Firth of Forth, a subset of the data presented by Wanless et al. (2018). The lengths are all standardised to 1 July, and they correspond well with simultaneous estimates from sandeel survey data (Wanless et al., 2004). Tuning effective handling time to this time series does not affect the predicted temporal trend or predicted relative differences in sandeel length between locations, only the absolute length.

Table S1. Model variables and parameter values, with sources where relevant. All parameters were rounded to two significant digits, unless this was not a sufficient level of precision in relation to the magnitude of uncertainty. Associated uncertainty ranges are included in brackets. d.l. = dimensionless.

| **Name** | **Description** | **Value** | **Unit** | **Source** |
| --- | --- | --- | --- | --- |
| *State variables* | | | | |
| $R$ | Reserve energy | - | kJ | - |
| $S$ | Structural energy | - | kJ | - |
| *Translated state variables* | | | | |
| $W$ | Wet weight sandeel | - | g | - |
| $L$ | Total length sandeel | - | cm | - |

| *Input variables* | | | | |
| --- | --- | --- | --- | --- |
| $T$ | Temperature | - | °C | Daily averages of temperature $T$ (°C) from the ERA5 Climate Reanalysis, which provides hourly sea surface temperature with a 31$\times$31 km resolution (Copernicus Climate Change Service C3S, 2017). |
| $\Sigma n_{k}$ | Food abundance for each prey type *k* | - | ind m^-3^ | Modelled abundances based on data collected by the Continuous Plankton Recorder (see Olin, 2020; Olin et al., 2022). |
| $E_{k}$ | Energy content of prey type *k*, see Table S2 | - | kJ | Sourced from the literature (see Olin, 2020; Olin et al., 2022). |
| $\psi_{k}$ | Image area of prey type *k*, see Table S2 | - | m^2^ | Sourced from the literature (see Olin, 2020; Olin et al., 2022). |
| $L_{k}$ | Length of prey type *k*, see Table S2 | - | mm | Sourced from the literature (see Olin, 2020; Olin et al., 2022). |
| $h_{day}$ | Hours of daylight | - | h | Sunrise to sunset, rounded to a whole number. Calculated using the function “daylength” in the R-package “geosphere” (Hijmans, 2017). |
| $I_{0}$ | Average surface irradiance during hours of daylight | - | W m^-2^ | Obtained from a Fortran subroutine sourced from Ljungström et al. (2020). $I_{0}$ is a function of cloud cover, which was assumed to be constant at 0.75 based on a rough estimate of the average value during summer in the study area (Giggenbach et al., 2010). |
| $a_{d}$ | Diffuse attenuation coefficient | - | m^-1^ | Based on observations from hydrodynamic regions in the North Sea corresponding to sandeel habitat (see supplementary materials in Capuzzo et al., 2018). |
| $a_{b}$ | Beam attenuation coefficient | - | m^-1^ | $a_{b}$ can be approximated by a linear relationship with  $a_{d}$ ($a_{b}$ = 5$a_{d}$ − 0:08; Shannon, 1975). |

| *Assimilated energy* | | | | |
| --- | --- | --- | --- | --- |
| $A$ | Assimilated energy | - | kJ day^-1^ | - |
| $h_{active}$ | Number of hours in a day available for feeding | - | h | - |
| $i_{h}$ | Energy intake during a given hour | - | kJ h^-1^ | - |
| $i_{max}$ | Maximum energy intake without gut space limitation | - | kJ h^-1^ | - |
| $\epsilon$ | Assimilation efficiency | - | d.l. | - |
| $p_{c}$ | Proportion of time spent in search class $c$ | - | d.l. | - |
| $\lambda_{k,L,I}$ | Search rate for prey type $k$, by sandeel length $L$, at light conditions $I$ | - | m^3^ h^-1^ | - |
| $D_{k,L,I}$ | Detection distance for prey type $k$, by sandeel length $L$, at light conditions $I$ | - | m | - |
| $\gamma_{L}$ | Visual sensitivity of a sandeel of length $L$ | - | d.l. | - |
| $I_{Z}$ | Ambient irradiance at feeding depth $z$ | - | W m^-2^ | - |
| $\phi_{k}$ | Capture probability of prey type $k$ | - | d.l. | - |
| $g$ | Gut content | - | g | - |
| $d$ | Digestion rate | - | g h^-1^ | - |
| $\delta_{food}$ | Energy density of ingested prey mix | - | kJ g wet weight^-1^ | - |
| $g_{max}$ | Maximum gut content | - | g | - |
| $\alpha_{\epsilon}$ | Faecal loss coefficient | 0.82 (0.73; 0.91) | d.l. | Based on experimental measurements on *A. dubius* (Larimer, 1992). Uncertainty ranges are based on measurements of minnow (*Phoxinus phoxinus*) (Cui & Wootton, 1988). |
| $\beta_{\epsilon}$ | Faecal loss temperature scaling factor | 7.6 (7.5; 7.7)✕10^-3^ | d.l. | Based on experimental measurements on *A. dubius* (Larimer, 1992). Uncertainty ranges are based on measurements of minnow (Cui & Wootton, 1988). |
| $U_{\epsilon}$ | Nitrogenous excretion loss coefficient | 5.1 (2.0; 12)✕10^-2^ | d.l. | Based on experimental measurements of minnow (Cui & Wootton, 1988), with the uncertainty range based on observations from a range of fish species (Elliott, 1976). |
| $h_{e}$ | Effective handling time | 50 (40; 60) | s | Tuned to time series of sandeel lengths from the Firth of Forth (see text above and Olin, 2020). Uncertainty range represents range of values that still provide plausible predictions. In our model, $h_{e}$ not only represents the time taken to capture and handle prey, but acts as a general restriction on feeding efficiency as a result of e.g. the sandeel having to spend time on other behaviours such as predator avoidance: see Olin (2020) for a more detailed discussion. |
| $v$ | Swimming speed | 1.5 (0.5; 2.0) | body lengths s^-1^ | Average speed based on experimental measurements of *A. tobianus* were used (van Deurs et al., 2010). Upper and lower boundaries were obtained from Christensen (2010). |
| $C$ | Prey contrast | 0.23 (0.20; 0.26) | d.l. | Based on measurements of transparent copepods at wavelengths comparable to those of *A. marinus* habitat (Utne-Palm, 1999). Range from measured standard deviations. |
| $D_{frac}$ | Fraction of $L$ equal to detection distance with no light limitation | 1/2 (1/3; 1) | d.l. | Based on experimental observations of foraging *A. marinus* (Christensen, 2010). Usually reacts at a distance of ⅓ (lower value) to ½ (nominal value) of its body length, at most one full body length (maximum value). |
| $K_{D}$ | Light saturation | 3.5 (2.0; 5.0) | μE m^-2^ s^-1^ | Experimentally estimated based on several species of fish (Aksnes & Utne, 1997), range given by the lowest and highest estimate, with the midpoint taken as the nominal value. |
| $z$ | Feeding depth | 30 (0; 70) | m | Nominal value taken from previous foraging model of *A. marinus* (van Deurs et al., 2015). As sandeels are found throughout the water column while foraging (Freeman et al., 2004; Johnsen et al., 2017), 0 m (i.e. the surface) was used as the minimum, and 70 m the maximum (upper depth limit of suitable habitat; Wright et al., 2000). |
| $b$ | Capture probability decline rate | 5.095 (4.075; 6.115) | d.l. | From relationship based on experiments on a range of fish species (Pepin et al., 1987), with uncertainty range representing the reported standard errors. |
| $m$ | Capture probability sigmoidal midpoint | -1.9 (-2; -1.8) | d.l. | Tuned to observed ratios of prey size in available and ingested prey (see text above and Olin, 2020). Uncertainty range represents range of values that still provide plausible predictions. |
| $\alpha_{dig}$ | Digestion coefficient | 3.5 (3.0; 4.0)✕10^-2^ | d.l. | Nominal value taken from previous foraging model of *A. marinus* (van Deurs et al., 2015), in turn based on observations of *A. tobianus* (van Deurs et al., 2010). Validated against data on *A. marinus* (see Olin, 2020; Figure S2). Range of uncertainty taken from estimates of the same parameters from other species of fish (Persson, 1979, 1981, 1982). |
| $\beta_{dig}$ | Digestion scaling factor | 5.4 (5.1; 5.7)✕10^-2^ | d.l. | See $\alpha_{dig}$. |
| $\delta*$ | Energy density reference prey | 4.4 (3.9; 4.9) | kJ g wet weight^-1^ | Nominal value taken from previous foraging model of *A. marinus* (van Deurs et al., 2015), uncertainty (standard deviation) from measurements from the same source as used by van Deurs et al. (2015) (Verkuil et al., 2006). |
| $\alpha_{gut}$ | Maximum gut size coefficient | 1.7 (1.6; 2.0)✕10^-3^ | d.l. | Estimated from North Sea *A. marinus* gut measurements collected April–October 2006–2010 (see e.g. van Deurs et al., 2014; N = 2143). Uncertainty represents the 95% confidence limit. |
| $\beta_{gut}$ | Maximum gut size scaling factor | 2.3 (2.2; 2.4) | d.l. | See $\alpha_{gut}$. |
| *Metabolic costs* | | | | |
| $M$ | Total daily metabolic costs | - | kJ day^-1^ | - |
| $M_{SMR}$ | Standard metabolic rate | - | kJ day^-1^ | - |
| $M_{feed}$ | Metabolic activity costs | - | kJ day^-1^ | - |
| $M_{SDA}$ | Metabolic costs from processing food | - | kJ day^-1^ | - |
| $\alpha_{met}$ | SMR coefficient | 4.5 (2.5; 6.5)✕10^-3^ | d.l. | Based on experimentally measured values in overwintering *A. marinus* (Wright et al., 2017; see calculations in Olin, 2020). Lower and upper values are based on the standard deviation. Corrected to summer acclimatisation using estimates from Quinn & Schneider (1991). |
| $\beta_{met}$ | Metabolic weight scaling factor | 0.65 (0.51; 0.79) | d.l. | Based on experimental measurements of summer-acclimatised *A. personatus* (Quinn & Schneider, 1991). Upper boundary from average measured value in fish, with a symmetrical lower boundary, which is still within reported values for fish (Clarke & Johnston, 1999). |
| $Q_{10}$ | Temperature effect on SMR | 3.1 (1.5; 3.4) | d.l. | Based on experimentally measured values in overwintering *A. marinus* (Wright et al., 2017). Lower boundary from overwintering *A. personatus* (Quinn & Schneider, 1991) and upper boundary from the maximum values observed in fish (Clarke & Johnston, 1999) |

| $F$ | Feeding costs | 3.4 (3.0; 3.8)✕10^-3^ | kJ g^-1^ h^-1^ | Estimated using experimental observations of *A. tobianus* (van Deurs et al., 2010). Uncertainty based on the standard deviation of estimates of feeding costs in capelin *Mallotus villosus* (Behrens et al., 2006). |
| --- | --- | --- | --- | --- |
| $\zeta_{SDA}$ | Cost of processing food | 0.16 (0.016; 0.59) | d.l. | Nominal value based on the average value from a large range of studies of fish, with lower and upper values based on the range of values observed (Secor, 2009). |
| *Allocation* | | | | |
| $f_{S}$ | Proportion of net energy gain allocated to structural energy | - | - | - |

| $\alpha_{alloc}$ | Allocation coefficient | 0.43 (0.42; 0.44) | d.l. | As structural energy by definition cannot decrease, the derivative of the relationship between total energy and structural energy in a population represents how structural energy increases as total energy increases, which is equivalent to the proportion allocated to structure. The relationship between structural energy and total energy was fit as a power function using length and weight data from *A. marinus* collected from foraging puffins on the Isle of May 1996–2015 from 1 June to 26 July (N = 155) (see e.g. Wanless et al., 2018), as well as from trawl and dredge surveys conducted 2000–2009 late May to June, also in the north-western North Sea (N = 6819) (see MacDonald, 2017). The uncertainty was based on the 95% confidence interval. The final parameters $\alpha_{alloc}$ and $\beta_{alloc}$ were then obtained by taking the derivative of the estimated parameters. |
| --- | --- | --- | --- | --- |
| $\beta_{alloc}$ | Allocation scaling factor | -0.09 (-0.10; -0.08) | d.l. | See $\alpha_{alloc}$. |

| *Translation* | | | | |
| --- | --- | --- | --- | --- |
| $\alpha_{dry}$ | Structural dry weight coefficient | 1.7 (1.6; 1.9) | d.l. | Estimated from dataset of *A. marinus* brought to the Isle of May by puffins (see e.g. Wanless et al., 2018) (N = 186). Ash, fat and protein dry weight, total length and wet weight were available for each individual. It was assumed that structural dry weight is made up of ash and non-remobilisable protein. Non-remobilisable protein as a function of length was obtained by assuming that those individuals with the lowest protein content for their length only contained non-remobilisable protein, fitting a log-log-relationship between length and protein to fall below the lowest protein content per length (see Olin, 2020 for details). Non-remobilisable protein was then predicted for each individual based on their length and added to the ash dry weight to obtain structural dry weight. $\alpha_{dry}$ and $\beta_{dry}$were then estimated by fitting a relationship between log10-transformed structural dry weight and log10-transformed length, with the 95% confidence intervals as a measure of uncertainty. |
| $\beta_{dry}$ | Structural dry weight scaling factor | 3.27 (3.22; 3.33) | d.l. | See $\alpha_{dry}$. |
| $\delta_{S}$ | Structural energy density | 19.2 (19.0; 19.4) | kJ g dry weight^-1^ | Based on the remobilisable protein predicted for each individual in the puffin dataset (as described above for $\alpha_{dry}$), structural energy $S$ was obtained by multiplying the non-remobilisable energy with the energy density of protein (23.7 kJ g^-1^; Crisp, 1971). Ash contains no energy. $S$ was then divided by the total dry weight (ash and non-remobilisable protein) to obtain $\delta_{S}$ for each individual and taking the mean of this, using the standard deviation to represent uncertainty. |
| $\delta_{R}$ | Reserve energy density | 27 (25; 29) | kJ g dry weight^-1^ | $R$ of each sandeel in the dataset was estimated by multiplying  the fat content of each sandeel in the puffin dataset (as described above for the parameter $\alpha_{dry}$) with the energy density of fat (39.6 kJ g^-1^; Crisp, 1971), and multiplying the remobilisable protein (total protein minus nonremobilisable  protein) with the energy density of protein (23.7 kJ g^-1^; Crisp, 1971) and adding the two together. $\delta_{R}$ was then estimated by dividing the estimated $R$ by the estimated reserve dry weight (fat dry weight plus remobilisable protein dry weight) for each sandeel and taking the mean. Again, uncertainty was represented by the standard deviation. |
| $\omega_{SDW}$ | Structural dry to wet weight conversion factor | 5.7 (4.7; 6.9) | d.l. | To estimate how both structural and reserve dry weights translate into wet weights ($\omega_{SDW}$ and $\omega_{RDW}$), the general approach by MacDonald et al. (2018) was followed, making use of a published linear relationship between the proportion of wet weight made up of fat and the proportion of wet weight made up of water in *A. marinus* (Hislop et al., 1991). As it is assumed that all fat is part of the reserve energy, the proportion of fat can be set to 0 in this equation in order to estimate the proportion of water in a sandeel made up completely of structural energy, from which $\omega_{SDW}$ can be estimated. The magnitude of uncertainty was assumed to be the same as for $\omega_{RDW}$ (see below). |
| $\omega_{RDW}$ | Reserve dry to wet weight conversion factor | 3.9 (3.2; 4.6) | d.l. | $\omega_{RDW}$ was estimated by calculating reserve wet weight for each each sandeel in the puffin dataset (as described above for the parameter $\alpha_{dry}$) by subtracting structural wet weight (which can be estimated using $\omega_{SDW}$ and the previously estimated structural dry weight) from total wet weight and dividing this by the sum of fat dry weight and previously estimated remobilisable protein dry weight (i.e. reserve dry weight) and taking the mean. The standard deviation was again used as a measure of uncertainty. |

To assess how our predictions depended on the choices of parameter values, we examined the difference in predicted lengths when (i) increasing and decreasing parameter values 10%, indicating which model processes have a particularly large impact on predictions, and when (ii) varying the parameter values based on lower and upper defined uncertainty boundaries. In brief, we found that model predictions were sensitive to parameters governing the maximum intake rate and the relationship between length and structural dry weight (Figure S1). Parameter values for the sandeel’s swimming speed, feeding depth, the sandeel’s visual acuity and the costs associated with processing food and synthesising tissue were the main sources of uncertainty in predictions.

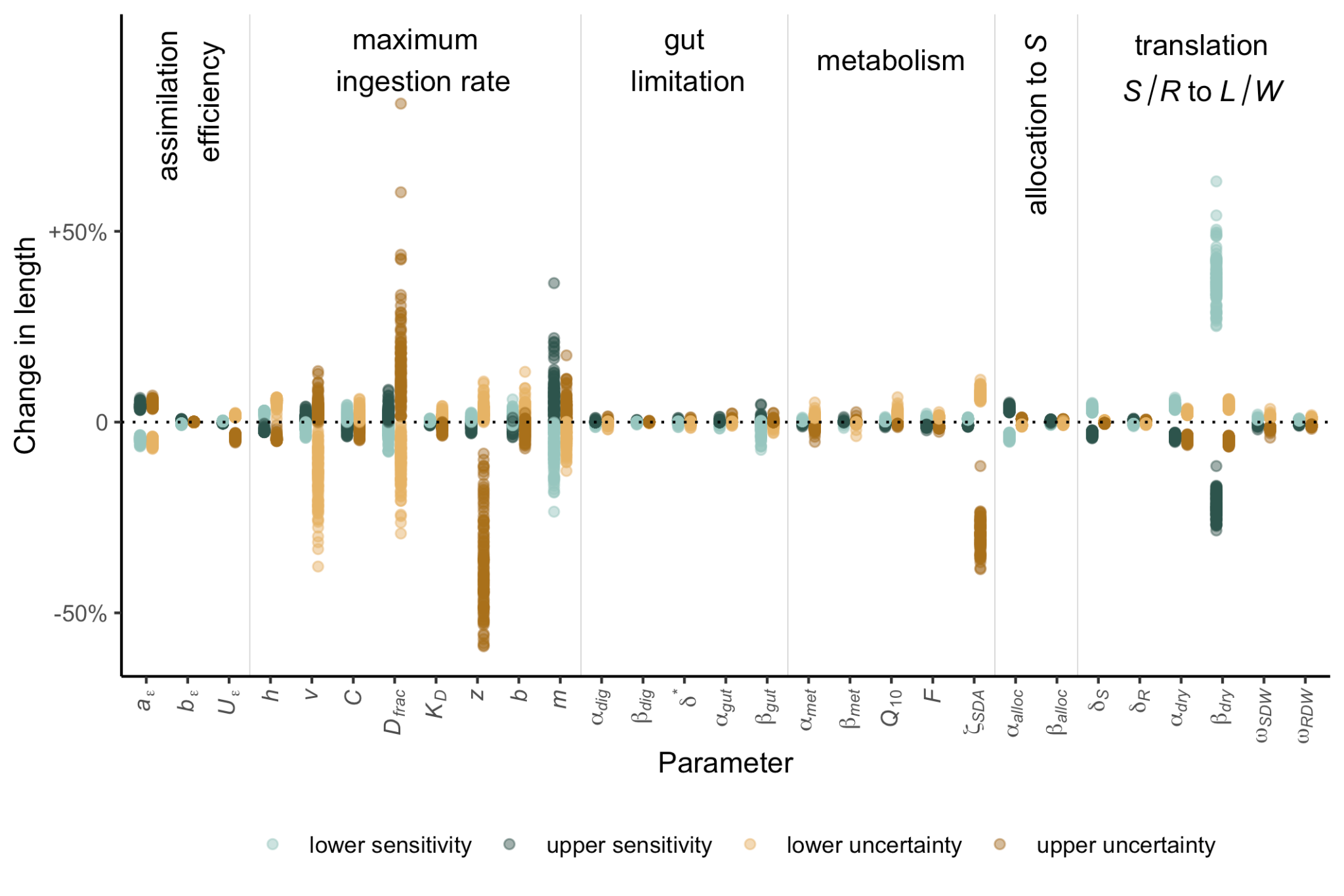

Figure S1. Parameter sensitivity analysis based on 10% decrease (light blue markers) and increase (dark blue markers), as well as the defined lower (light brown markers) and upper (dark-brown markers) uncertainty boundaries, with each point within each category representing a different location-year. y-axis shows predicted length at time of overwintering in relation to the baseline scenario of all parameters at their nominal value. For the meaning of each parameter, see Table S1. The general process each parameter belongs to is indicated at the top.

### SI3: Assimilated energy

Table S2. Trait values for the prey types included in the model. For sources of information on length, energy content and image area see Olin et al. (2022). For the search classes, these were divided into small copepods <1.3 mm and other small prey <1.3 mm (A), large copepods >1.3 mm (B), and large crustaceans and fish larvae (C) (see SI3.7).

| **Species** | **Length**  **(mm)** | **Energy content (kJ)** | **Image area**  **(mm^2^)** | **Search class** |
| --- | --- | --- | --- | --- |
| *Acartia* spp. | 1.15 | 0.385 | 0.29 | A |
| Appendicularia | 1 | 0.2537 | 0.39 | A |
| *Calanus finmarchicus* | 2.7 | 2.816 | 1.61 | B |
| *Calanus helgolandicus* | 2.68 | 2.772 | 1.59 | B |
| *Calanus* I–IV | 1.65 | 0.874 | 0.60 | B |
| *Calanus* V–VI | 2.48 | 2.376 | 1.36 | B |
| *Centropages hamatus* | 1.3 | 0.476 | 0.37 | B |
| *Centropages typicus* | 1.63 | 0.77 | 0.59 | B |
| *Centropages* spp. | 1.55 | 0.72 | 0.53 | B |
| Copepod nauplii | 0.19 | 0.00435 | 0.028 | A |
| Decapoda larvae | 0.9 | 1.472 | 0.64 | A |
| Euphausiacea spp. | 17 | 252 | 28.37 | C |
| *Evadne* spp. | 0.5 | 0.0924 | 0.20 | A |
| Fish eggs | 1 | 2.72 | 0.79 | A |
| Fish larvae | 12 | 4 | 6.57 | C |
| Hyperiidea spp. | 16 | 92.8 | 54.84 | C |
| *Metridia lucens* | 2.27 | 1.395 | 1.14 | B |
| *Oithona* spp. | 0.68 | 0.09 | 0.10 | A |
| *Para-Pseudocalanus* | 0.7 | 0.1292 | 0.11 | A |
| *Podon* spp. | 1 | 0.924 | 0.79 | A |
| *Temora longicornis* | 1 | 0.24 | 0.22 | A |

##### SI3.1 Assimilation efficiency

Faecal losses have been estimated in the closely related *A. dubius* and are positively related to temperature (Gilman, 1994; based on measurements from Larimer, 1992). However, there are no measurements of nitrogenous excretion in any species of sandeels, and it is unknown how it responds to for example temperature (which is often found to have a positive impact on excretion rates, see e.g. Elliott, 1976). As such, it is assumed that a constant proportion $U_{\epsilon}$ of ingested energy is lost to nitrogenous excretion. Combining the two types of losses, the following equation for $\epsilon$ is obtained:

$$\epsilon=(\alpha_{\epsilon}+\beta_{\epsilon}T)-U_{\epsilon}$$

(S3)

where $T$ is temperature (°C), and $\alpha_{\epsilon}$, $\beta_{\epsilon}$ and $U_{\epsilon}$ are constants. See Table S1 for how parameter values were sourced.

##### SI3.2 Maximum intake rate and optimal foraging

The processes included in the calculated maximum intake rate are described in the main text. Maximum intake $i_{max}$ of given prey search class $c$ is given by:

$$i_{max,c}=\frac{\sum_{k} \lambda_{k,L,I}\phi_{k}n_{k}E_{k}}{1+\sum_{k} h_{e}\lambda_{k,L,I}\phi_{k}n_{k}}$$

(S4)

where $\lambda_{k,L,I}$ is the search rate for prey type $k$ by a sandeel of length $L$ at light conditions $I$ (m^3^ h^-1^; see SI3.3, SI3.4, SI3.5), $\phi_{k}$ is the capture rate for prey type $k$ (see SI3.6), $n_{k}$ is the prey density (individuals per m^3^; see “Environmental data” in the main text), $E_{k}$ is the energy content per individual prey (kJ; see Table SI2) and $h_{e}$ is the handling time (s). Three different prey search classes were considered based on experimental and field observations (Christensen, 2010; Eigaard et al., 2014; Godiksen et al., 2006; van Deurs et al., 2014): small copepods and other small prey (<1.3 mm; prey class A), (ii) large copepods (>1.3 mm; prey class B) and (iii) large crustaceans and fish larvae (prey class C) (see SI3.7, Table S2 and Olin, 2020). It is assumed that the sandeels only forage on one search class at the time, and that the time spent in each search class is proportional to its profitability (Visser & Fiksen, 2013).

The proportion of time spent in each search class is thus calculated as follows:

$$p_{C}=\frac{i_{max, c}}{\sum_{c} i_{max,c}}$$

(S5)

where $i_{max, c}$ is the maximum potential energy ingested during one hour spent feeding in search class $c$ (see Eq. S4). As digestion is modelled as a separate process, differences in digestion rates are not included when determining profitability. The evidence for digestive quality (i.e. the amount of energy that can be assimilated per unit of digestion time) as a basis for prey selection is mixed (Fall & Fiksen, 2020; Gill & Hart, 1998; Kaiser et al., 1992), and as sandeel stomach fullness is often well below maximum capacity (Olin, 2020), digestive quality likely plays a minor role in prey selection. While it is possible that the energy expenditure may vary depending on feeding mode (see e.g. Queiros et al., 2024), this has not been studied in sandeels and is therefore not included in the model.

To obtain the amount of energy that can be ingested within an hour $i_{max}$ (kJ h^-1^) the energy consumed in each prey class is added up:

$$i_{max}=\sum_{c} p_{c}i_{max,c}$$

(S6)

where $p_{c}$ is the proportion of time spent in each search class (Eq. S5), and $i_{max,c}$ (kJ h^-1^) is the maximum amount of energy that can be ingested in a given hour in a given search class, assuming continuous feeding and ignoring the limitation from gut space (Eq. S4).

##### SI3.3 Search rate

The search rate $\lambda_{k,L,I}$ (m^3^ h^-1^) is a function of the prey size-, sandeel size- and light-dependent detection distance $D_{k,L,I}$ (m; see SI3.4) and the length of the sandeel $L$, as this determines speed:

$\lambda_{k,L,I}=D_{k,L,I}^{2}\pi vL$

(S7)

where $D_{k,L,I}$ (m) is the detection distance, $v$ is the swimming speed (body lengths per hour) and $L$ (m) is the sandeel length. $vL$ provides the speed in m h^-1^. The equation assumes a cylindrical search space with a cross-section of ${D_{k,L,I}}^{2}\pi$ where the radius is equal to the detection distance (thus assuming that the angle of the visual field is 90°). By multiplying this with the swimming speed, the volume searched per hour (m^3^ h^-1^) is obtained. This is a common model of planktivorous fish foraging (e.g. Eggers, 1977), and is also used in the van Deurs et al. (2015) model.

##### SI3.4 Detection distance

The detection distance $D_{k,L,I}$ (m) is, as in van Deurs et al. (2015), dependent on prey size, sandeel size and light conditions. It is based on the model of visual range in fish developed by Aksnes & Utne (1997), and modelled as follows:

$$D_{k,L,I}^{2}\exp\left( a_{b}D_{k,L,I} \right)=C\psi_{k}\gamma_{L}\frac{I_{z}}{K_{D}+I_{z}}$$

(S8)

where $a_{b}$ is the beam attenuation coefficient (input to the model, dependent on turbidity, see the main text and Table S1), $C$ is the prey contrast, $\psi_{k}$ (m^2^) is the image area of a given prey type (Table S2), $\gamma_{L}$ is the sandeel length-dependent visual sensitivity of sandeels (see below), $K_{D}$ (μE m^-2^ s^-1^) the light saturation of detection distance and $I_{z}$ (W m^-2^) the ambient irradiance (see SI3.5). The equation is solved through iteration, based on an adaptation of the Fortran code presented in the supplementary materials of Ljungström et al. (2020).

Visual sensitivity $\gamma_{L}$ is calculated based on the distance at which prey is detected when there is no light limitation (i.e. $I_{z}/(K_{D}+I_{z}) =1$). Here, this was done using experimental observations by Christensen (2010) of foraging sandeels under well-lit conditions. It is assumed that visual sensitivity scales with body size, as is generally observed in planktivorous fish (e.g. Miller et al., 1993). Visual sensitivity $\gamma_{L}$ is thus calculated as follows:

$$\gamma_{L}=\frac{{(D_{frac}L)}^{2}}{C}$$

(S9)

where $L$ (m) is sandeel length, $D_{frac}L$ (m) is the measured detection distance (thus assuming detection distance at no light limitation scales with body length and is a constant fraction of the body length), $C$ is the prey contrast and $\psi_{ref}$ (m^2^) is the prey image area of the experimental prey used when measuring detection distances, here that of the 7 mm herring larvae used in the experiment.

##### SI3.5 Ambient irradiance $I_{Z}$

The ambient irradiance $I_{Z}$ (W m^-2^) is calculated as follows:

$I_{z}=I_{0}e^{-a_{d}z}$

(S10)

where $I_{0}$ (W m^-2^) i is the surface irradiance on a given day (average irradiance during the foraging period, which is input to the model), $a_{d}$ (m^-1^) is the diffuse attenuation coefficient (also model input, dependent on turbidity) and $z$ (m) is the foraging depth.

##### SI3.6 Capture probability $\phi_{k}$

In planktivorous fish, it has repeatedly been found that larger prey items are more difficult to capture (Folkvord & Hunter, 1986; Fuiman, 1989; Margulies, 1989). This likely contributes to the finding that the average size of prey in sandeel guts is only slightly greater than the average size of prey in the water column (Godiksen et al., 2006), in spite of greater encounter rates for large prey types (see van Deurs et al., 2015 and previous sections) and potentially also an active preference for larger prey (Christensen, 2010). To account for the reduced capture success of larger prey types, capture probability is a function of prey size in the model. This relationship is often found to be nonlinear (Folkvord & Hunter, 1986; Fuiman, 1989), generally following a sigmoidal relationship (Pepin et al., 1987), which we also adopted here:

$\phi_{k}=(\frac{1}{1+exp(-b(ln(L_{k}/10-m)))})$

(S11)

where $L_{K}$is the size (mm) of prey type $k$ and $b$ and $m$ are constants. This results in a close-to-guaranteed success when targeting small prey, and then a sigmoidal decline as prey becomes larger. $b$ controls the steepness of the decline whereas $m$ controls the point at which the decline occurs.

##### SI3.7 Search classes

Each prey type also needs to be assigned to a search class, which captures the assumption that sandeels will only focus on one search class at a time, spending time in each search class in proportion to its profitability (Visser & Fiksen, 2013). Within each search class, prey types are similar and it would be expected that the sandeels use a common search image for these prey types. While behaviour, such as swimming speed, is likely to vary between search classes (see e.g. Queiros et al., 2024), not enough information is available to incorporate this and it is assumed that behaviour is the same for all search classes.

To delineate the different search classes, previous studies of *A. marinus* were used. Observations of foraging sandeels that were first feeding on *Acartia* spp., which have a length of around 1.15 mm (Richardson et al., 2006), found that the sandeels switched to feeding solely on herring (*Clupea harengus*) larvae (7mm) when these were introduced, while completely ignoring the copepods (Christensen, 2010). As such, these prey types are part of different search classes. This is supported by Eigaard et al. (2014) finding that sandeel larvae (minimum length 12 mm) and copepods (no reported length) were found in separate, distinct clumps in sandeel guts sampled in the field (suggesting that they only fed on one at a time). Further, Godiksen et al. (2006) found that krill (6 mm) and capelin larvae (18.3 mm) were clumped together, and suggested that this may be because the sandeel develop a common search image based on the larger size and darkly pigmented eyes. Copepods (2.5 mm) occurred in separate clumps. This suggests that fish larvae and large crustaceans are part of the same search class, and that large copepods are part of a separate search class. Finally, van Deurs et al. (2014) found that individual sandeel stomachs tended to contain either copepods smaller than 1.3 mm or copepods larger than 1.3 mm, suggesting that these size groups belong to different search classes. Together, this suggests three prey search classes: small (<1.3 mm) copepods (search class A), large (>1.3 mm) copepods (search class B) and fish larvae as well as other large crustaceans (search class C). Other small prey types such as cladocerans (all smaller than 1.3 mm) were grouped with the small copepods (search class A). The search class assigned to each prey type can be found in Table S2.

##### SI3.8 Maximum gut size

Based on the finding that the maximum gut size $g_{max}$ (g) in fish is well described by a power function of length (Pirhonen et al., 2019) it was modelled as follows:

$$g_{max}=\alpha_{gut}L^{\beta_{gut}}$$

(S12)

where $L$(cm) is sandeel length and $\alpha_{gut}$ and $\beta_{gut}$ are constants.

##### SI3.9 Digestion

Based on observations in *Ammodytes* spp. (Mackinson, 2007; Sun et al., 2010; van Deurs et al., 2010), and as is generally the case in fish (e.g. Andersen, 1999; Elliott, 1991), digestion is modelled as an exponential process. In addition, as is again observed both in *Ammodytes* spp. (van Deurs et al., 2010) and other species of fish (e.g. Andersen, 1999; Elliott, 1991), the rate increases with temperature. Finally, prey energy density is a key control on digestion rates in fish (Andersen, 1999, 2001), with the rate decreasing with increased energy density. This was included in the foraging model of *A. marinus* by van Deurs et al. (2015), where digestion rates were assumed to be inversely proportional to the energy density of the prey. We adopt the same equation as van Deurs et al. (2015), modelling digestion rate $d$ (g h^-1^) as:

$d=\alpha_{dig}exp(\beta_{dig}T)\frac{\delta*}{\delta_{food}}g$

(S13)

where $\alpha_{dig}$ and $\beta_{dig}$are constants, $T$ (°C) is temperature, $\delta*$ (kJ g wet weight^-1^) is the energy density of the prey used when experimentally measuring digestion rates, $\delta_{food}$ (kJ g wet weight^-1^) is the overall energy density of prey in the gut and $g$ is the gut content (g).

There are published observations on gut evacuation in *A. marinus* (Mackinson, 2007), but as the temperature and prey energy density were not reported, this information could not be used for parameterisation, but was instead used for validating the chosen parameters. The observations were based on sandeels caught in the south-western North Sea in June and then kept in a tank, with 10 sandeels sampled each hour to measure gut contents. For input, the temperature for mid-June for the centre of the sampling area in Mackinson (2007) (based on data from Copernicus Climate Change Service C3S 2017) and prey energy density for the typical sandeel diet in this area around this time (van Deurs et al., 2013) were used. This resulted in a good agreement with observations (Figure S2a). Observations of *A. personatus* feeding on adult and nauplii of *Artemia salina* (from Sun et al., 2010) were also compared to model predictions. Reported energy densities, temperature, initial stomach content and sandeel size from Sun et al. (2010) were used as input. This also seemed to suggest that predictions were similar to observations, but less so for the nauplii diet (Figure S2b).

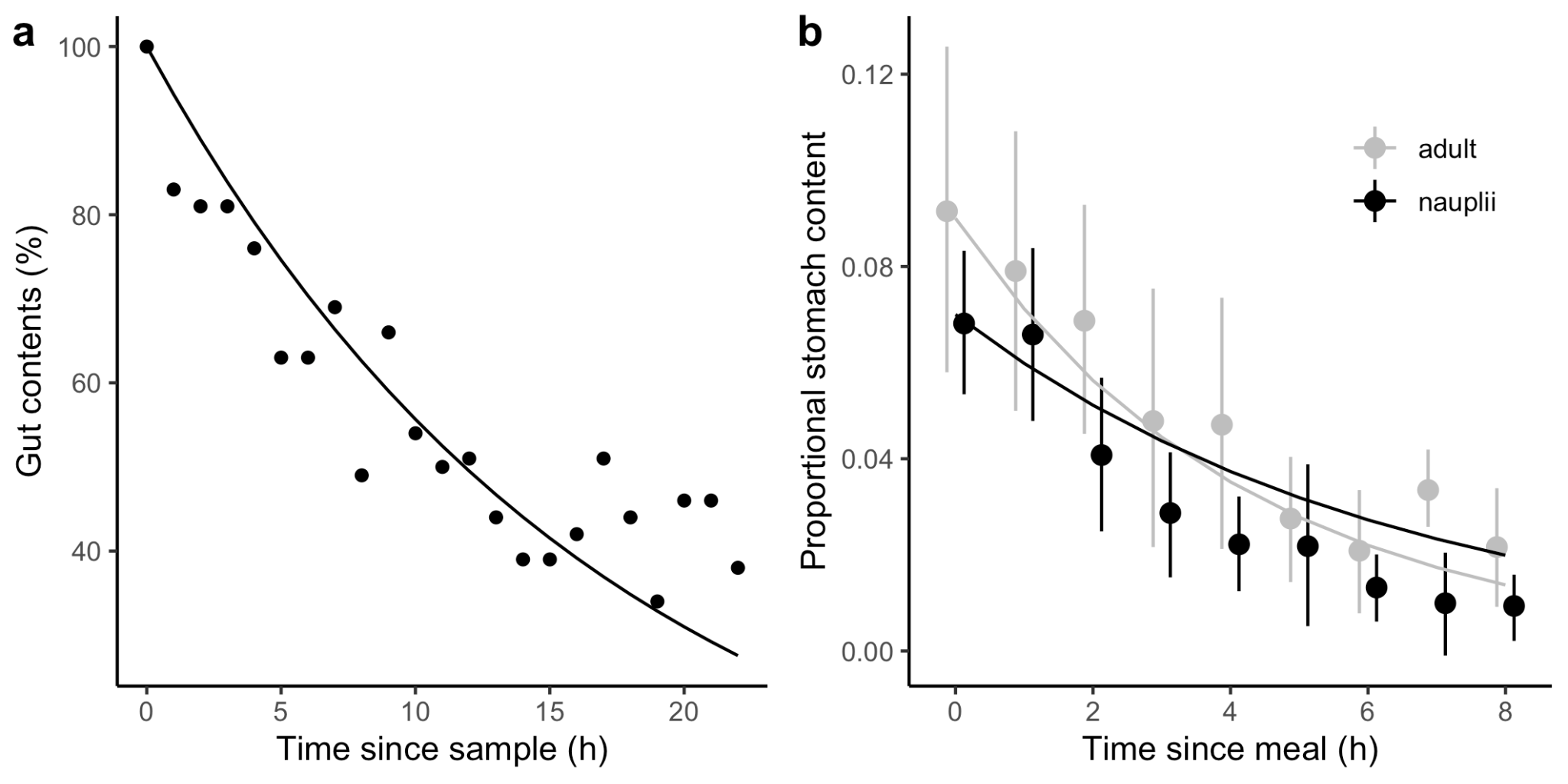

Figure S2 (a) Model predictions as compared to measurements from *A. marinus* (Mackinson, 2007). Markers show relative stomach content at sequential intervals following sampling in the field. Lines show corresponding model predictions based on Eq. S13. (b) Model predictions as compared to measurements from *A. personatus* (Sun et al., 2010). Markers with associated error bars are taken from estimates of relative gut content (wet weight of food divided by wet weight of sandeel) at sequential intervals following a meal, feeding on adults (grey) and nauplii (black) of *Artemia salina*. Lines show corresponding model predictions based on Eq. S13.

It is assumed that digestion occurs continuously, both during the feeding period and during the night. Anything left in the gut after a night of burrowing is carried through to the next feeding period, acting as a limitation on intake. This is based on experimental observations of *A. tobianus* showing that when food was very abundant, there was generally food left in the gut the next morning (van Deurs et al., 2011). In the field, guts collected first thing in the morning have been found to be empty (Wright & Bailey, 1993), but this may be expected as food is likely to be

more scarce than in the experimental setting.

##### SI3.10 Gut fullness and realised intake rate $i_{h}$

Finite gut space puts an ultimate limit on the amount of food a sandeel can ingest, and will depend on the rate of ingestion and digestion, as well as maximum gut capacity ($g_{max}$), which was assumed to be a function of sandeel size (see SI3.8). Digestion $d$ (g h^-1^), which removes food from the stomach at an hourly rate (i.e. reducing $g$, the current stomach content in grams), was modelled as a function of temperature and the energy density of the ingested food (see Andersen, 1999, 2001; Elliott, 1991; Mackinson, 2007; Sun et al., 2010; van Deurs et al., 2010; SI3.9).

Gut content is tracked in terms of wet weight and it is then assumed that intake cannot be greater than what fills the remaining space. It is assumed that weight and volume are proportional based on water content of zooplankton generally being high (Kiørboe, 2013). Gut content $g$ (g) thus changes as follows:

$\frac{dg}{dt}=\frac{i_{h}}{\delta_{food}}-d$

(S14)

where gut content increases with ingested food $i_{h}$ (kJ h^-1^) – as this is measured in terms of energy, it is divided by the energy density of the food $\delta_{food}$ (kJ g wet weight^-1^) to obtain ingested food in terms of wet weight — and decreases with digested food $d$ (g h^-1^). In the model, changes in gut content are discretised assuming hourly time steps.

The maximum intake in an hour is then corrected for available gut space to obtain the amount of food ingested in a given hour:

$i_{h}=\left\{ \begin{aligned} i_{max}, &g_{max}-g)\delta_{food}\geq i_{max} \\ \left( g_{max}-g \right)\delta_{food}, &g_{max}-g)\delta_{food}<i_{max} \end{aligned} \right.$

(S15)

where $i_{max}$ (kJ) is the maximum amount of food that can be ingested assuming no gut limitation (see SI3.2), $g_{max}$(g) is the maximum amount of food that can be held in the stomach, $g$ is the amount of food currently in the stomach (g, see Eq. S14) and $\delta_{food}$ is the energy density of the ingested prey mix (kJ g wet weight^-1^).

### SI4: Metabolic rate

#### SMR

The standard metabolic rate $M_{SMR}$ (kJ day^-1^) in fish is generally found to vary with body size and temperature (Clarke & Johnston, 1999), and so it was assumed that in sandeels it also takes on the commonly observed relationship:

$M_{SMR}=\alpha_{met}W^{\beta_{met}}Q_{10}^{T/10}$

(S16)

where $\alpha_{met}$ is a constant, $W$ (g) is sandeel wet weight, $\beta_{met}$ is the metabolic weight scaling exponent, $T$ (°C) is temperature and $Q_{10}$ represents the rate of change as temperature increases by 10 °C.

#### Activity costs

Determining the cost of activity is difficult and often a large source of uncertainty in bioenergetic modelling (Ney, 1993). The main determinants of activity costs in fish include swimming speed and fish mass as well as to some degree, temperature (Brett & Groves, 1979; Jobling, 1993). Here only the effect of swimming speed and fish mass were included, given the wide range of relationships between active metabolic rate and temperature observed in fish (see Fig. 2 in Brett, 1965) and in absence of any information on similar species. As such, $M_{feed}$ (kJ day^-1^) was be estimated as follows:

$$M_{feed}=FWh_{day}$$

(S17)

#### where $F$ (kJ g^-1^ h^-1^) is the cost of feeding per gram of fish per hour, $W$ (g) is sandeel wet weight, and $h_{day}$ (h) is the total number of hours of daylight. It was thus assumed that the costs are paid both when feeding ($h_{active}$) as well as during the two hours used for ascent and descent.

#### Specific dynamic action

The most important driver of variation in specific dynamic action (SDA) is the amount and type of food ingested (Secor, 2009). No relevant observations of SDA exist for any species closely related to *A. marinus.* For this reason, the simplifying assumption was made that SDA is proportional to the energy content of the meal (which seems to work relatively well in fish, Secor, 2009), so that $M_{SDA}$ (kJ day^-1^) is calculated as follows:

$$M_{SDA}= \zeta_{SDA}\frac{A}{\epsilon}$$

(S18)

where $A$ (kJ day^-1^) is the assimilated energy, is the assimilation efficiency (where $A/\epsilon$ = is equal to total ingested energy during the day, before accounting for assimilation efficiency), and $\zeta_{SDA}$ is the SDA-coefficient.

### SI5: Energy allocation

It is assumed that the allocation strategy changes after reserves have been depleted to prioritising rebuilding reserves (see main text): Based on this, the proportion $f_{S}$ of net energy gain allocated to structural energy was modelled as follows:

$$f_{S}=\left\{ \begin{aligned} \alpha_{alloc}(R+S)^{\beta_{alloc}}, R+\left( A-M \right)>R_{ideal} \\ 0, &R+\left( A-M \right)\leq R_{ideal} \end{aligned} \right.$$

(S19)

where $\alpha_{alloc}$ and $\beta_{alloc}$ govern how allocation to structure decreases as the total energy content (kJ) of the sandeel ($R+S$) increase, and $R+(A-M)\leq R_{ideal}$ indicate that when adding net energy gain for one day $(A-M)$ to reserves $R$ does not bring reserves up to the ideal level $R_{ideal}$ given the structural energy (or, length) of the sandeel, everything is allocated to reserves.

The idea that the relationship between $S$ and $R+S$ can be interpreted as the ideal ratio between $S$ and $R+S$ was used to estimate$R_{ideal}$. $R_{ideal}$(kJ) can thus be calculated as follows:

$$R_{ideal}=\left( \frac{S(\beta_{alloc}+1)}{\alpha_{alloc}} \right)^{1/(\beta_{alloc}+1)}-S$$

(S20)

where $\alpha_{alloc}$ and $\beta_{alloc}$ are constants and $S$ (kJ) is structural energy.

Note that if adding the net assimilated energy $A-M$ to reserves $R$ brings reserves up to above

pre-starvation levels, some allocation to structural energy will occur before body composition is restored. This fits with previous observations of post-starvation allocation in fish (see review in Jones, 2001).

### SI6: Results

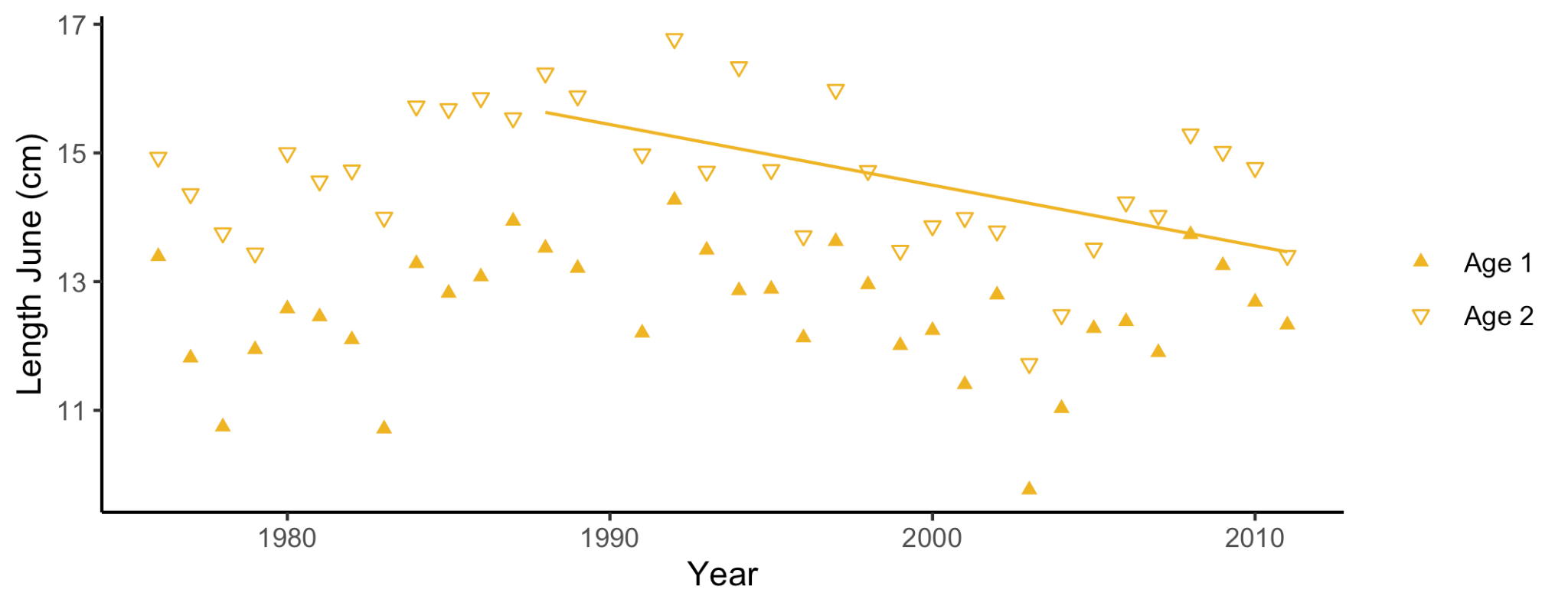

Figure S3. Lengths of age 1 and age 2 sandeels in Dogger Bank based on fisheries data from June (extracted from van Deurs et al., 2014). The length of age 2 sandeels declined 1988–2011 with 0.09 cm per year (95 % CI: -0.16; -0.02; linear regression) but the confidence intervals of the decline in age 1 sandeels clearly overlapped with 0 [estimate -0.05; 95 % CI: -0.11; 0.01; linear regression]).

Table S3. Relationships between predicted length at overwintering and average temperature during the growth season, and between observed lengths and average temperatures up until the time of observation. Estimates show the estimated parameters for a linear model (LM) and for the two terms of a polynomial model (PM), with 95 % CIs. The final column shows ΔAIC_C_ of the linear model and the polynomial model, both compared to a null model. If a model had ΔAIC_C_ ≥ 4 compared to a simpler model, we considered the relationship supported (highlighted in bold). Corresponding data are shown in Figure 4b−c.

|  | Location | Estimates [95 % CI] | Likelihood ratio test |
| --- | --- | --- | --- |
| Predicted length vs. temperature | Shetland  N = 36 | LM: 0.47 [-0.0044; 0.94]  PM^1^: -9.6 [-22; 2.3]  PM^2^: 0.47 [-0.091; 1.0] | ΔAIC_C_ linear = -1.7  ΔAIC_C_ polynomial = -2.5 |
|  | ECG  N = 23 | LM: -0.22 [-0.80; 0.36]  PM^1^: -5.5 [-21; 10]  PM^2^: 0.21 [-0.40; 0.82] | ΔAIC_C_ linear = -3.1  ΔAIC_C_ polynomial = -6.5 |
|  | Firth of Forth  N = 23 | LM: -0.17 [-0.83; 0.49]  PM^1^: -1.9 [-18; 14]  PM^2^: 0.07 [-0.61; 0.75] | ΔAIC_C_ linear = -3.1  ΔAIC_C_ polynomial = -6.6 |
|  | Dogger Bank  N = 33 | LM: 0.24 [-0.027; 0.52]  PM^1^: -3.5 [-12; 4.7]  PM^2^: 0.15 [-0.18; 0.47] | ΔAIC_C_ linear = -1.7  ΔAIC_C_ polynomial = -5.6 |

| Observed length vs. temperature | Hermaness  N = 11 | LM: -0.96 [-2.8; 0.88]  PM^1^: -6.2 [-69; 57]  PM^2^: 0.27 [-3.0; 3.6] | ΔAIC_C_ linear = -2.6  ΔAIC_C_ polynomial = -7.1 |
| --- | --- | --- | --- |
|  | Fair Isle  N = 24 | LM: -1.6 [-2.2; -0.92]  PM^1^: -11 [-34; 11]  PM^2^: 0.51 [-0.64; 1.7] | **ΔAIC_C_ linear = 6.0**  ΔAIC_C_ polynomial = 4.2 |
|  | ECG  N = 6 | LM: -0.027 [-1.4; 1.4]  PM^1^: 53 [18; 88]  PM^2^: -1.9 [-3.2; -0.65] | ΔAIC_C_ linear = -29  ΔAIC_C_ polynomial = too few observations to compute |
|  | Firth of Forth  N = 22 | LM: -0.018 [-0.43; 0.39]  PM^1^: -1.8 [-11; 7.3]  PM^2^: 0.079 [-0.34; 0.50] | ΔAIC_C_ linear = -4.3  ΔAIC_C_ polynomial = -8.9 |
|  | Dogger Bank  N = 13 | LM: 0.24 [-0.34; 0.83]  PM^1^: 0.48 [-46; 47]  PM^2^: -0.0088 [-1.7; 1.7] | ΔAIC_C_ linear = -4.3  ΔAIC_C_ polynomial = -8.3 |

Table S4. Relationships between predicted length at overwintering and the examined prey field characteristics during the growth season, and between observed lengths and examined prey field characteristics up until the time of observation. Estimates show the estimated parameters for a linear model (LM) and for a log-model (LOGM), with 95 % CIs. The final column shows ΔAIC_C_ of the linear model and the log-model, both compared to a null model. If a model had ΔAIC_C_ ≥ 4 compared to a simpler model, we considered the relationship to be supported (highlighted in bold). Corresponding data are shown in Figure 5b−c.

|  | Location | Estimates [95 % CI] | Likelihood ratio test |
| --- | --- | --- | --- |
| Predicted length vs. total energy | Shetland  N = 36 | LM: 0.00030 [0.00021; 0.00039]  LOGM: 2.0 [1.4; 2.7] | ΔAIC_C_ linear = 7.8  **ΔAIC_C_ log = 21** |
|  | ECG  N = 23 | LM: 0.00029 [0.00011; 0.00048]  LOGM: 2.8 [1.5; 4.0] | ΔAIC_C_ linear = -11  **ΔAIC_C_ log = 12** |
|  | Firth of Forth  N = 23 | LM: 0.00023 [-0.000027; 0.00050]  LOGM: 1.3 [0.19; 2.4] | ΔAIC_C_ linear = -16  ΔAIC_C_ log = 2.5 |
|  | Dogger Bank  N = 33 | LM: 0.000096 [-0.00012; 0.00031]  LOGM: 0.61 [-1.0; 2.3] | ΔAIC_C_ linear = -18  ΔAIC_C_ log = -1.1 |

| Predicted length vs. *Calanus finmarchicus* | Shetland  N = 36 | LM: 0.0065 [0.0034; 0.0096]  LOGM: 0.68 [0.34; 1.0] | ΔAIC_C_ linear = -0.17  **ΔAIC_C_ log = 8.9** |
| --- | --- | --- | --- |
|  | ECG  N = 23 | LM: 0.0029 [0.0014; 0.0045]  LOGM: 1.5 [0.87; 2.1] | ΔAIC_C_ linear = -4.0  **ΔAIC_C_ log = 13** |
|  | Firth of Forth  N = 23 | LM: -0.0068 [-0.035; 0.021]  LOGM: -0.46 [-1.2; 0.25] | ΔAIC_C_ linear = -9.4  ΔAIC_C_ log = -1.6 |
|  | Dogger Bank  N = 33 | LM: -0.036 [-0.070; -0.0028]  LOGM: -0.60 [-1.7; 0.47] | ΔAIC_C_ linear = -5.2  ΔAIC_C_ log = -1.3 |
| Predicted length vs. prey size | Shetland  N = 36 | LM: 0.55 [-1.9; 3.0]  LOGM: 1.2 [-4.0; 6.5] | ΔAIC_C_ linear = -0.048  ΔAIC_C_ log = 1.5 |
|  | ECG  N = 23 | LM: 2.1 [0.46; 3.8]  LOGM: 5.3 [1.2; 9.5] | **ΔAIC_C_ linear = 4.0**  ΔAIC_C_ log = 5.9 |
|  | Firth of Forth  N = 23 | LM: -3.0 [-5.1; -1.0]  LOGM: -7.0 [-12; -1.9] | **ΔAIC_C_ linear = 6.5**  ΔAIC_C_ log = 7.2 |
|  | Dogger Bank  N = 33 | LM: 3.1 [-2.2; 8.5]  LOGM: 6.2 [-4.4; 17] | ΔAIC_C_ linear = 2.1  ΔAIC_C_ log = 3.4 |

| Observed length vs. total energy | Hermaness  N = 11 | LM: 0.00011 [-0.00027; 0.00049]  LOGM: 1.0 [-2.8; 4.8] | ΔAIC_C_ linear = -20  ΔAIC_C_ log = -1.8 |
| --- | --- | --- | --- |
|  | Fair Isle  N = 24 | LM: -0.000065 [-0.00029; 0.00016]  LOGM: -0.50 [-2.1; 1.1] | ΔAIC_C_ linear = -19  ΔAIC_C_ log = -1.1 |
|  | ECG  N = 6 | LM: 0.00013 [-0.00016; 0.00042]  LOGM: 1.1 [-0.23; 2.5] | ΔAIC_C_ linear = -45  ΔAIC_C_ log = -27 |
|  | Firth of Forth  N = 22 | LM: 0.00002 [-0.00017; 0.00021]  LOGM: 0.20 [-0.71; 1.1] | ΔAIC_C_ linear = -20  ΔAIC_C_ log = -2.6 |
|  | Dogger Bank  N = 13 | LM: 0.00068 [0.00042; 0.00094]  LOGM: 2.5 [1.2; 3.8] | ΔAIC_C_ linear = -7.9  **ΔAIC_C_ log = 5.4** |
| Observed length vs. *Calanus finmarchicus* | Hermaness  N = 11 | LM: -0.0036 [-0.014; 0.0067]  LOGM: -0.40 [-1.3; 0.48] | ΔAIC_C_ linear = -14  ΔAIC_C_ log = -4.2 |
|  | Fair Isle  N = 24 | LM: 0.0015 [-0.0027; 0.0057]  LOGM: 0.30 [-0.26; 0.85] | ΔAIC_C_ linear = -13  ΔAIC_C_ log = -2.6 |
|  | ECG  N = 6 | LM: 0.0046 [0.0025; 0.0067]  LOGM: 1.2 [0.65; 1.8] | ΔAIC_C_ linear = -33  ΔAIC_C_ log = -23 |
|  | Firth of Forth  N = 22 | LM: -0.0083 [-0.045; 0.028]  LOGM: -0.046 [-0.46; 0.36] | ΔAIC_C_ linear = -9.0  ΔAIC_C_ log = -4.3 |
|  | Dogger Bank  N = 13 | LM: 0.0037 [-0.016; 0.023]  LOGM: 0.20 [-0.45; 0.84] | ΔAIC_C_ linear = -12  ΔAIC_C_ log = -4.4 |
| Observed length vs. prey size | Hermaness  N = 11 | LM: 8.2 [4.2; 12]  LOGM: 15.8 [8.1; 23] | **ΔAIC_C_ linear = 8.1**  ΔAIC_C_ log = 9.6 |
|  | Fair Isle  N = 24 | LM: 1.6 [-1.5; 4.7]  LOGM: 3.1 [-3.7; 9.8] | ΔAIC_C_ linear = 0.73  ΔAIC_C_ log = 2.1 |
|  | ECG  N = 6 | LM: 2.7 [-1.6; 7.0]  LOGM: 6.5 [-3.9; 17] | ΔAIC_C_ linear = -25  ΔAIC_C_ log = -24 |
|  | Firth of Forth  N = 22 | LM: 0.031 [-1.0; 1.1]  LOGM: 0.083 [-2.6; 2.8] | ΔAIC_C_ linear = -2.5  ΔAIC_C_ log = -0.57 |
|  | Dogger Bank  N = 13 | LM: 1.1 [-2.1; 4.3]  LOGM: 2.4 [-4.0; 8.9] | ΔAIC_C_ linear = -1.1  ΔAIC_C_ log = 0.40 |
